# DNA Damage Driven Viability Loss and Transcriptional Reprogramming in Chinese Hamster Ovary Cell Perfusion Culture

**DOI:** 10.64898/2026.03.16.712126

**Authors:** Noah B Hitchcock, Maxine Annoh, Luigi Grassi, Samik Das, Cristina Sayago Ferreira, Daniel Ray, Ramy Elgendy, Lin Wang, Ken Lee, Ian M Sudbery, Daniel A Bose, Diane Hatton, Si Nga Sou, Rajesh K Mistry, Christopher P Toseland

**Affiliations:** Drug Substance Development, BioPharmaceutical Development, BioPharmaceuticals R&D, AstraZeneca, Cambridge, CB2 0AA, UK; Division of Clinical Medicine, School of Medicine and Population Health, University of Sheffield, Sheffield, S10 2RX, UK; Drug Substance Development, BioPharmaceutical Development, BioPharmaceuticals R&D, AstraZeneca, Gaithersburg, USA; Analytical Sciences, BioPharmaceutical Development, BioPharmaceuticals R&D, AstraZeneca, Cambridge, CB2 0AA, UK; Translational Genomics, Discovery Sciences, BioPharmaceuticals R&D, AstraZeneca, Gothenburg, Sweden; Central Laser Facility, Research Complex at Harwell, Science and Technology Facilities Council, Rutherford Appleton Laboratory, Harwell, Didcot, Oxford, OX11 0QX, UK; School of Biosciences, Faculty of Science, University of Sheffield, Sheffield, S10 2TN, UK

## Abstract

Intensified perfusion cultures promise higher yields and consistent product quality, yet extended runs frequently stall due to declining Chinese hamster ovary (CHO) cell viability at high cell density. Here, we identify DNA damage accumulation as a central, previously underappreciated driver of this limitation. Using two antibody-expressing CHO cell lines operated at a control and high cell density in perfusion bioreactors, we combined time-resolved transcriptomics with molecular, biophysical and super-resolution imaging analyses. We observed a progressive, global downregulation of DNA damage response (DDR) pathways accompanied by a time-dependent accumulation of DNA lesions. Notably, γH2AX signalling declined despite DNA damage, indicating impaired damage sensing and repair. Concomitantly, RNA polymerase II protein levels and transcriptional hub organisation were markedly reduced, consistent with widespread transcriptional dysfunction preceding loss of viability. An extended 21-day perfusion run confirmed continued viability decline beyond day 14, supporting a cumulative damage model. Comparison with HEK293 cells revealed substantially more efficient DNA repair, underscoring an intrinsically attenuated DDR in CHO cells. Together, these findings demonstrate that unrepaired DNA damage fundamentally constrains perfusion culture longevity, exposing limits of CHO genomic plasticity and highlighting DDR pathways as promising targets for host cell engineering to enhance perfusion performance.

## INTRODUCTION

Chinese hamster ovary (CHO) cells represent the predominant mammalian host cell line for the industrial manufacture of protein-based biotherapeutics because of their suitability for suspension culture and large-scale up in bioreactors, their capacity to generate human-like post-translational modifications, and their inherent genetic tractability and adaptability (1, 2). The ability of CHO cells to adapt to environmental changes has facilitated the implementation of intensified bioreactor production modes (3), such as continuous perfusion culture, in which cellular production phases are extended by continuously supplying fresh medium and feed whilst simultaneously removing product for downstream processing (1, 4, 5), thereby enabling superior product quality and yield (6). Hence, the biopharmaceutical sector has begun to pivot towards perfusion bioreactors as its preferred platform in order to leverage these benefits.

Perfusion processes are commonly described in two phases: a growth phase and a production phase **(Figure 1A)**. The growth phase encompasses the period during which the perfusion culture increases in viable cell density (VCD) until reaching a predefined target. Once this target is achieved, the process transitions into the production phase, during which the culture is maintained at the desired VCD. The continuous supply of fresh medium provides a favourable growth environment that can drive VCD beyond the intended set point. To prevent exceeding this threshold, a bleed feature is applied, whereby a defined volume of culture is removed to reduce the VCD and stabilise the culture at the target cell density (7). The duration of the growth phase depends on bioreactor seeding density, with higher seeding densities shortening the time to reach the target VCD (8), as well as reducing the length of the production phase due to declining viability **(Supplementary Figure S1A)**. However, when lower seeding densities are used to enable extended runs, the maintenance of the production phase at high viability remains limited to ∼10 days. The implementation of perfusion culture has been hampered by limitations with achieving the high VCDs and long-term CHO cell survival required to maximise productivity gains. Complexities arise due to process-related constraints within the bioreactor such as suboptimal gas transfer (9) and shear stress (10, 11), as well as cellular stressors including oxidative stress (3) and hypoxia (12). These stressors are hypothesised to intensify at elevated VCD and are thought to contribute to declining viability during attempts to extend bioreactor runs to ≥30 days. Indeed, studies reporting prolonged operations often achieve this only at reduced cell densities (13–15), highlighting a trade-off between run duration and VCD (16–21) **(Supplementary Figure S1A),** or alternatively rely on engineered cell lines to mitigate these limitations (16).

**Figure 1.**
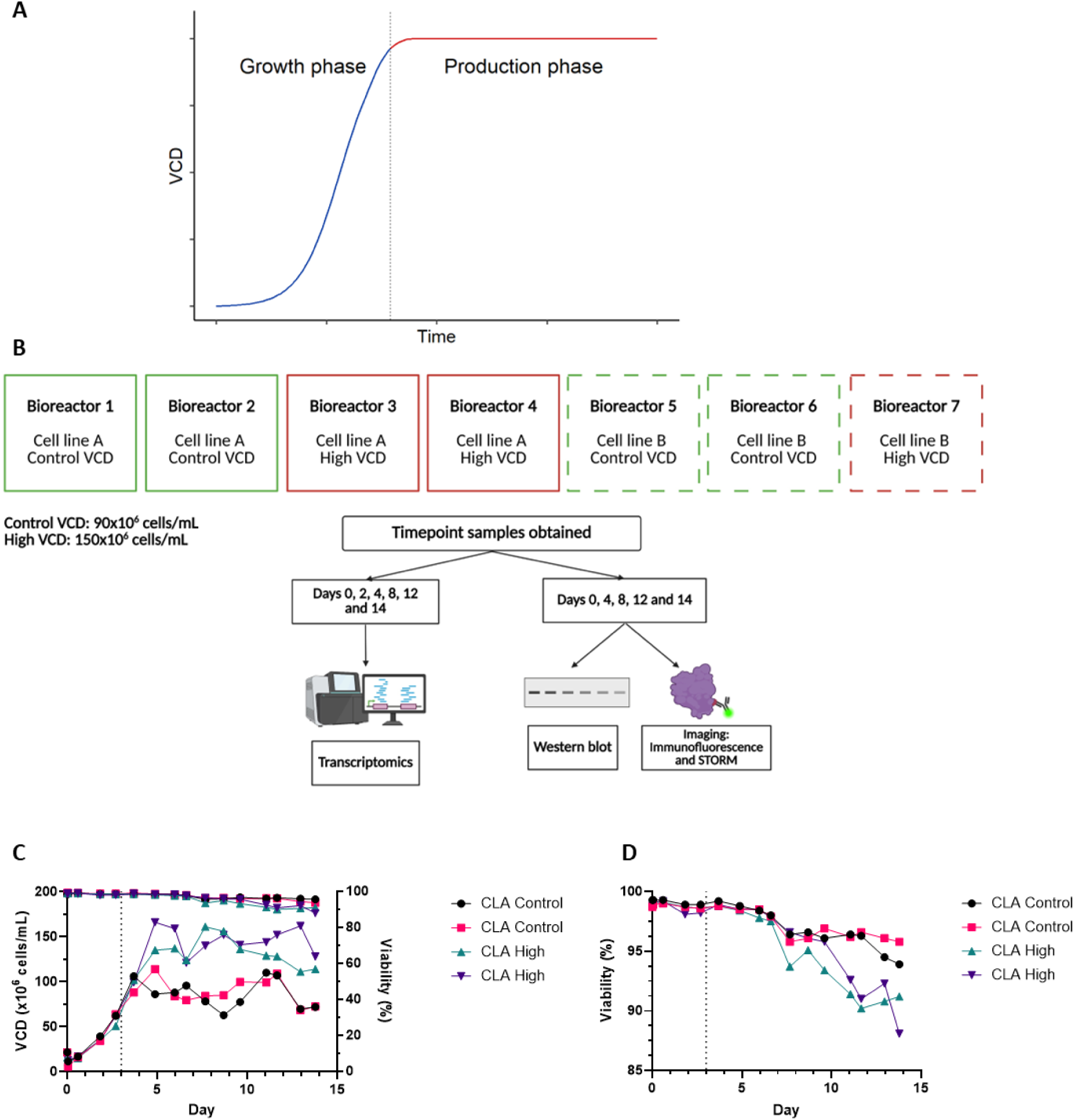
Perfusion bioreactor setup and growth kinetics. (A) Schematic depicting a typical perfusion process growth profile displaying viable cell density (VCD) over time. The blue and red line represent the growth and production phases, respectively. The vertical grey line separates the two phases. (B) The perfusion bioreactor setup used in this investigation. Solid and dashed outlines depict cell line A (CLA) and cell line B (CLB), respectively. Green outlines show control VCD target (90×10^6^ cells/mL) and red outlines show high VCD target (150×10^6^ cells/mL). Timepoints sampled for transcriptomics, western blot and immunofluorescence imaging are denoted. Two biological replicates were performed per condition and one biological replicate for cell line B high VCD. (C) VCD (left y-axis) and viability (right y-axis) over the 14-day perfusion bioreactor run. Dashed line at day 3 indicates the start of the maintenance of the target VCD. (D) Viability on a shortened axis to highlight the high VCD bioreactors lower end of run viability.

A key feature of CHO cells is their marked genomic instability (22) and plasticity (2, 23, 24), which is thought to stem, in part, from an attenuated DNA damage response (DDR), including known mutations in DDR genes in CHO-K1 derived lineages (22, 25). While such plasticity has been exploited for host evolution (26–29) and performance improvements (3), its consequences under continuous perfusion conditions have not been systematically evaluated. Here, we address this gap by phenotypically characterising control, and high VCD perfusion bioreactor runs across two independent antibody-expressing CHO cell lines, integrating transcriptomic, molecular, and biophysical profiling. Our results reveal progressive DNA damage accumulation as a potential driver of viability decline, accompanied by global downregulation of DDR pathways and insufficient DNA lesion repair. This is associated with genome-wide transcriptional reprogramming, reductions in transcription hub number (nuclear clustering of RNA polymerase II molecules), and altered spatial organisation indicative of impending transcriptional collapse. Collectively, these findings position perfusion culture as a high-stress environment that exposes the limits of CHO cell genomic plasticity, identifying DNA damage and transcriptional reprogramming as critical bottlenecks that limit perfusion duration and highlight actionable targets to improve performance.

## MATERIALS AND METHODS

### Cell Lines and Culture Conditions

AstraZeneca proprietary CHO-K1 derived host cells were maintained in CD-CHO medium (Life Technologies) supplemented with 6 mM L-glutamine (Life Technologies) and sub-cultured every 3-4 days at a seeding density of 0.2–0.3×10^6^ cells/mL. Human embryonic kidney (HEK) FreeStyle™ 293-F Cells (Gibco™) were maintained in CD 293 media supplemented with 6 mM L-glutamine (Life Technologies) and sub-cultured every 3-4 days at a seeding density of 0.2-0.3×10^6^ cells/mL. Stably transfected CHO cell pools and cell lines producing an easy-to-express (ETE) monoclonal antibody (mAb) were routinely maintained in AstraZeneca proprietary medium. Routine culture for all cell lines was performed at 36.5°C in vented, non-baffled Erlenmeyer flasks (Corning, Amsterdam, The Netherlands), in 5 % CO_2_ maintained at 140 rpm. Cells were sub-cultured every 3-4 days at a seeding density of 0.3-0.4×10^6^ cells/mL. Viable cell density (VCD) and cell viability were assessed using a Vi-Cell XR or BLU automated cell counter and viability analyser by trypan blue exclusion (Beckman Coulter, High Wycombe, UK).

### Ambr 15 Fed-Batch Cell Line Screen

Cell lines derived from stable pools (30) were inoculated into Ambr 15 cell culture bioreactors (Sartorius AG, Göttingen, Germany) with a 1×10^6^ cells/mL seeding density in 15 mL of production medium. Bolus additions of AstraZeneca production feeds were supplemented over the 14-day Ambr run. pH was measured using an ABL90 FLEX PLUS (Radiometer Medical ApS, Brønshøj, Denmark) and adjusted with the addition of base to maintain pH 7. The dissolved oxygen (DO_2_) and CO_2_ were maintained at 50% and 0%, respectively. Glucose and lactate were measured using a YSI 2900D Biochemistry Analyser (YSI Inc, Yellow Springs, OH, USA). Bolus glucose additions were supplemented to maintain an 8 g/L culture concentration. Titre measurements of mAb were obtained by protein-A HPLC affinity chromatography on an Agilent 1260 Infinity series (Agilent Technologies, CA, USA) by comparing the peak size from each sample with a calibration curve. Specific cell productivity (qP) was calculated as follows: qP = Th/CCTf (where Th is the Harvest Titre and CCTf is the calculated Cumulative Cell Time on the last day of the culture). CCTi = ((di − di-1) × (VCNi + VCNi-1)/2) + CCTi-1 (where d is the day of the culture, VCN is the viable cell count and i is the day of VCN sampling during the course of the culture).

### Gene Copy Analysis by Droplet Digital PCR

Genomic DNA was extracted from 1×10^7^ cells using a PureLink ™ Genomic DNA kit (Invitrogen, CA, USA) according to the manufacturer’s protocol. Gene copy number analysis was carried out on Not1-digested genomic DNA from each sample using a TaqMan Assay System with probes designed against antibody encoding DNA sequence (Thermo Fisher Scientific, MA, USA) and a Q×100 Droplet Digital PCR System, along with QuantaSoft software (BioRad, CA, USA). Manufacturer’s instructions were followed throughout.

### Measurement of mRNA Levels

Cellular RNA was extracted from 1×10^7^ cells using an RNAeasy mini plus kit (Qiagen, Hilden, Germany) according to the manufacturer’s protocol. RNA concentration was measured using a NanoDrop Eight (Thermo Fisher Scientific, MA, USA). 3 µg RNA was reverse transcribed to generate cDNA using a SuperScript IV First-Strand Synthesis System (Thermofisher Scientific, MA, USA) according to the manufacturer’s instructions. Quantitative (q)PCR reactions were conducted in 20 µL reactions using 20x TaqMan Assay Probes against antibody encoding DNA sequence. The *Mmadhc* gene was used as the reference (31). qPCR reactions were performed using a QuantStudio 12K Flex Real-Time PCR System (Applied Biosciences, CA, USA). The ΔCt method was used to calculate the mRNA levels of the HC and LC transcripts.

### Flow Cytometry

Cell lines were collected by centrifugation, fixed in 70% methanol, and stored at −20 °C before further processing. CHO host cells were used as unstained and stained controls. For HC and LC staining, cells were washed twice on ice with DPBS containing 1% BSA, followed by a centrifugation step at 130 x g for 5 min to remove the washing solution. For staining, 2 mL 1% BSA prepared in DPBS supplemented with 5 µg/mL APC-conjugated goat anti-human IgG Fcγ (Jackson ImmunoResearch, Cat. No. 109–136–170) and 3 µg/mL Kappa-FITC goat anti-human antibody (Southern Biotechnology, Cat. No. 2060–02) were added to each sample. Samples were then incubated in the dark on ice for 60 min before they were washed twice with 1% BSA. Finally, cell pellets were resuspended in 1 mL 1% BSA in DPBS and 200 μL of each sample were transferred into a Nunc™ 96-well polypropylene microplate (ThermoFisher Scientific, CA, USA, Cat. No. 249944). The stained cells were analysed by a MACSQuant flow cytometer with the APC and FITC double-positive population measured according to manufacturer’s specifications. Data analysis was performed using FlowJo (BD, NJ, USA) software. Perfusion Bioreactor Two independent perfusion studies were conducted: an initial 14-day run using seven bioreactors (Cell line A: two control VCD target vessels and two high VCD target vessels; Cell line B: two control VCD target vessels and one high VCD target vessel) and an extended 21-day run using four bioreactors (Cell line A only: two control VCD target vessels and two high VCD target vessels). Cells were cultured in perfusion mode using 3 L glass stirred tank bioreactors (Chemglass Life Sciences, Vineland, NJ) equipped with a tangential flow filtration (TFF) system for cell retention. The hollow fibre cartridge (S04-P20U-10-N, Repligen, Waltham, MA) enabled continuous perfusion throughout the culture period. Bioreactors were controlled via DASGIP software (Eppendorf, Hamburg, Germany) throughout. Scale-up vessels were inoculated at a 0.8×10⁶ cells/mL seeding density before the perfusion studies to ensure adequate growth rate. To initiate the perfusion bioreactor run, the cultures were inoculated at 10×10⁶ cells/mL. A vessel volume per day (VVD) of 1.2 was maintained throughout. Culture bleeding was implemented on day 3 to prevent exceeding the pre-determined target VCD setpoints of 90×10⁶ cells/mL (control) or 150×10⁶ cells/mL (high), controlled by a predictive algorithm. Permeate flow rates were adjusted accordingly to maintain constant VVD. Process parameters were maintained at 35.5°C, 50% dissolved oxygen, and pH 7.0 ± 0.10 (controlled via CO₂ sparging). Agitation started at 300 rpm and increased in response to dissolved oxygen controller output. AstraZeneca proprietary medium and feed was used throughout, and glucose was kept at 3.3 g/L. Daily samples were analysed for pH, glucose, lactate, pO₂, and pCO₂ using a RAPIDPoint 500e analyser (Siemens, Malvern, PA). Cell counts and protein titre were measured as stated previously.

### RNA Sample Preparation and Transcriptomic Sequencing

Samples for RNA-sequencing were taken on days 0, 2, 4, 8, 12 and 14. At least 1×10^7^ cells were collected from each bioreactor and spun at 130 g for 5 min, supernatant removed, and pellet resuspended in 200 μL RNAlater® (Sigma-Aldrich, CA, USA, Cat. No. R0901), snap frozen on dry ice and stored at −80°C before further processing. Upon removal of RNAlater, cell pellets were treated with lysis buffer mix containing proteinase K sourced from an RNAdvance Tissue kit (Beckman Coulter). Cells were mixed and subsequently incubated at 37°C for 25 min. RNA extraction was performed according to the manufacturer’s protocol on a Biomek i7 Hybrid robotic workstation (Beckman Coulter). Subsequent steps involved mRNA enrichment and sequencing library preparation utilising a KAPA mRNA HyperPrep Kit (Roche), according to the manufacturer’s instructions, using a Tecan Fluent® liquid handler. Library integrity and quality were evaluated using a SS NGS fragment kit (1–6000 bp; Agilent) on a fragment analyser. Sequencing was performed as paired-end 2 × 150 reads on an Illumina NovaSeq6000 platform, using a 300-cycle S1 Reagent Kit v1.5 (Illumina).

### Transcriptomic Analysis

Sequence reads were aligned to the proprietary vector sequence reference, using bwa mem (version 0.7.17) (32, 33) with default parameters. Samtools view (version 1.11) (34) was used with the option “-f 12” to identify read pairs where both mates were non mapped to the proprietary vector reference. These unmapped read pairs were subsequently used for all downstream analyses. To trim the sequencing and PCR adapters, fastp (version 0.23.4) (35) was used with the options “--compression 9 --detect_adapter_for_pe --qualified_quality_phred 15 --unqualified_percent_limit 10 --average_qual 20 --length_required 30”. Transcript expression was quantified using trimmed and qc filtered reads with Salmon (version 1.10.0) (36), with the selective alignment strategy (37) and the parameters “--libType A --validateMappings – mimicBT2”, using annotations from Ensembl CHOK1GS (version 93) (38). The Bioconductor package txtimport (37) was used to summarise transcript expression into gene expression (39, 40). The raw RNA-sequencing counts were normalised using variance stabilising transformation (VST) from DESeq2 v1.42.1 (41) for principal component analysis (PCA) and network analysis. Weighted gene co-expression network analysis (WGCNA) (42) was conducted using the gene whole co-expression network analysis (GWENA) R package v1.16.0 (43). The signed network was built using VST normalised counts calculated by Pearson correlation coefficient. Modules containing highly co-expressed genes were detected from the network and 0.9 threshold was used to merge modules with similar co-expression patterns. Functional enrichment of modules were calculated using the bioconductor enrichR (Version 3.2) library (https://cran.r-project.org/web/packages/enrichR/) to query the Enrichr database (44) Differential gene expression analysis was mase using DESeq2 (Version 1.42.1). Comparisons between high and control VCD bioreactors were conducted separately for each day using two modelling approaches: a likelihood ratio test (LRT) that accounted for cell line differences, and an alternative model that excluded the cell line term. Genes were considered differentially expressed if they had an adjusted p-value <0.01 and log_2_ fold change >0.58. Gene Set Enrichment Analysis (GSEA) was performed using clusterProfiler (Version 3.2), (45) with genes ranked by log_2_ fold change.

### Western Blot Sample Preparation

Samples for western blot were taken on days 0, 4, 8, 12 and 14. At least 1×10^7^ cells were collected from each bioreactor and spun at 4000 rpm for 10 mins, supernatant removed, snap frozen on dry ice and stored at −80°C before further processing.

### Western Blot

Cell pellets were lysed using a buffer containing 50 mM Tris pH 7.0 (Thermo Fisher Scientific, MA, USA), 2% UltraPure SDS Solution (Thermo Fisher Scientific, MA, USA) and boiled at 80°C. The lysed pellets were sonicated in preparation for quantification. The protein was quantified using a Pierce Dilution-Free Rapid Gold BCA Protein Assay (Thermo Fisher Scientific, MA, USA) following the manufacture’s protocol. A standard curve of BSA (Thermo Fisher Scientific, MA, USA) was prepared and Abs_480_ was measured on a SpectraMaxID plate reader (Molecular Devices, Wokingham, UK). 20 μg protein was prepared with NuPAGE LDS Sample Buffer (Thermo Fisher Scientific, MA, USA) and run on a NuPAGE Bis-Tris Midi Protein gel, 4-12%, 1.0 mm (Thermo Fisher Scientific, MA, USA) with either the PageRuler or PageRuler Plus Prestained Protein Ladder (Thermo Fisher Scientific, MA, USA). Transfer was conducted using the iBlot 2 Dry Blotting System (Thermo Fisher Scientific, MA, USA) and a PVDF iBlot Transfer Stack (Thermo Fisher Scientific, Waltham, MA, USA). The transfer conditions were as follows, 20 V for 1 min, 23 V for 4 min and 25 V for 5 min. The membrane was washed with 1X TBS-Tween (Thermo Fisher Scientific, MA, USA) and blocked with 5% BSA in TBS-Tween for 1 h at room temperature. The membrane was washed and incubated with 1:1000 primary antibody **(Supplementary Table 1)** in 5% BSA TBS-Tween at 4°C overnight. GAPDH was used as loading control. The membrane was washed and incubated with IRDye 800 CW goat anti-rabbit IgG secondary antibody (LI-COR bio, NE, USA) and IRDye 680RD goat anti-mouse IgG secondary antibody (LI-COR bio, NE, USA) at 1:10000 in 5% BSA in TBS-Tween for 45 min at room temperature. The membrane was developed and imaged using an Odyssey CLx Imager (LI-COR bio, NE, USA). Densitometry was performed using Fiji ImageJ v1.53. Supplementary Table 1 contains further information on the antibodies used.

### Immunofluorescence, Comet Assay, Atomic Force Microscopy and STORM Sample Preparation

Samples for immunofluorescence imaging, STORM or comet assays were taken on days 0, 4, 8, 12 and 14. At least 1×10^7^ cells were collected from each bioreactor and spun at 130 g for 5 min, supernatant removed, resuspended in proprietary AstraZeneca medium supplemented with 10% DMSO (Sigma-Aldrich, CA, USA) and stored at −80°C before further processing. All frozen samples were processed identically to preserve any proportional differences between samples.

### Immunofluorescence

The frozen cell aliquots were revived and seeded on poly-D-lysine (Sigma-Aldrich, CA, USA) coated glass coverslips No 1.5 12 mm diameter. Cell nuclei were stained with Hoechst 33342 (Thermo Fisher Scientific, MA, USA) and incubated for 10 min at 37°C. The stain was removed and washed with PBS and fixed in 4% *w/v* PFA in PBS. Residual PFA was quenched with 50 mM NH_4_Cl (Thermo Fisher Scientific, MA, USA) in PBS for 15 min. The cells were permeabilised and blocked with 2% w/v BSA (Sigma-Aldrich, CA, USA), 0.1% *v/v* triton X-100 (Sigma-Aldrich, Burlington, CA, USA) in TBS. RNAPII-pSer5 was stained with rabbit mAb to RNA polymerase II CTD repeat YSPTSPS Alexa Fluor 488 (Phospho S5) (1:500, Ab196152; Abcam, Cambridge, UK) with 1% BSA in TBS and incubated for 1 h at room temperature. γH2AX was stained with rabbit P-Histone H2A.X(S139) antibody (1:200, Cat: 9718, Cell Signaling Technology, MA, USA) for 1 h at room temperature. The coverslips were washed and incubated with the secondary conjugated antibody, donkey anti-rabbit 488 secondary antibody (1:500, ab181346; Abcam, Cambridge, UK) with 1% BSA in TBS and incubated for 1 h. RNAPII-pSer2 was stained with rabbit mAb to RNA polymerase II CTD repeat YSPTSPS Alexa Fluor 488 (Phospho S2) (1:200, Ab237279; Abcam, Cambridge, UK) with 1% BSA in TBS and incubated at 4°C overnight. The coverslips were subsequently washed with TBS and H_2_O, then mounted onto microscope slides with Mowiol (Sigma-Aldrich, CA, USA) supplemented with 2.5% w/v DABCO (Sigma-Aldrich, Burlington, CA, USA). Supplementary Table 1 contains further information on the antibodies used.

### Confocal Microscopy

Fixed cells were imaged using a Zeiss LSM980, with a Plan-Achromat 63 × 1.4 NA oil immersion objective (Carl Zeiss, 420782-9900-000). Two laser lines: 405 and 488 nm were used for excitation of Hoechst and Alexa-fluor 488 fluorophores. A built-in multi-band dichroic mirror MBS405/488/561 (Carl Zeiss, 1784-995) was used to reflect excitation laser beams onto samples. For fluorescence signal collection, the used emission spectral bands were: 410–524 nm (Hoechst) and 493–578 nm (Alexa-fluor 488). The green channel (Alexa-fluor 488) was imaged using a 1 gallium arsenide phosphide (GaAsP) detector, while the blue (Hoechst) channels were imaged using two multi-anode photomultiplier tubes (MA-PMTs). For imaging acquisition and rendering, Zeiss ZEN Blue software (v2.3) was used.

### Widefield Imaging

Widefield images were acquired using a Zeiss Cell Discoverer 7 with a 20x 0.7 NA Plan-Apochromat air objective. LED illumination (385nm and 469nm) corresponding to Hoechst and Alexa-Fluor488 were used with multi bandpass filter sets. Images were captured with a Hamamatsu Orca Flash 4.0 V3 camera using the Zeiss ZEN Blue software (v2.3). Nuclear shape and fluorescence intensities were measured subtracting background fluorescence using Fiji ImageJ v1.53 software.

### Alkaline Comet Assay

To detect both single, double-stranded breaks and alkali labile sites in DNA, a Comet assay kit (ab238544, Abcam, Cambridge, UK) was used following an adapted version of the manufacturer’s protocol. CHO cells were revived then washed in ice cold PBS and resuspended to 1×10^6^ cells/mL. The cells were mixed at a 1:10 *v/*v with 37°C comet agarose and dispensed onto the comet slides. The slides were immersed in lysis buffer at 4°C for 90 min followed by unwinding alkaline solution at 4°C for 90 min. The slides underwent electrophoresis in a horizontal gel tank at 18 V and 300 mA for 30 min. The cells were washed in cold deionised water and fixed in cold 70% ethanol. After fixing the cells, the slides were left to air dry at room temperature and stained with Vista Green DNA dye. The cells were imaged on a ZEISS LSM980 using a 10x/0.45 objective with a WD of 2 mm under an FITC filter. Comet tails were analysed using CometScore 2.0.

### Atomic Force Microscopy (AFM)

CHO cells were revived and seeded on a WillCo-dish pre-treated with poly-D-lysine (Sigma-Aldrich, CA, USA) and allowed to adhere for 30 min. AFM measurements were performed with a Bruker Bioscope Resolve mounted on an inverted microscope (Nikon Eclipse Ti2) connected to an ORCA-Flash4.0LT (Hamamatsu) camera. Imaging was performed in filtered PBS at room temperature. Pre-calibrated Silicon Tip – Nitride cantilevers (PFQNM-LC-V2 Bruker) were used with a 70 nm tip radius. PeakForce Tapping/PeakForce Capture with a maximum indentation force of 0.5 nN was used. Indentation curves were fitted within Nanoscope Analysis (Bruker) using a cone-sphere model (46).

### Stochastic Optical Reconstruction Microscopy (STORM)

STORM measurements followed established protocols (47, 48). Round 25 mm No 1.5 coverslips were treated with the etching solution prepared with 5:1:1 distilled water, NH_4_OH (Sigma-Aldrich, CA, USA) and H_2_O_2_ (Thermo Fisher Scientific, MA, USA) at 70°C for 1 h. The coverslips were subsequently washed in methanol followed by distilled water then allowed to air dry. The coverslips were exposed to UV for 15 min before poly-D-lysine treatment. Frozen cell samples were thawed and 1×10^6^ cells were immediately dispensed onto the pre-treated poly-D-lysine coverslips. Cells were left to adhere for 30 mins at 37°C and fixed for 15 min in 4% *w/v* PFA in PBS. The residual PFA was quenched with 50 mM ammonium chloride in PBS for 15 min. Cells were permeabilised and blocked in 3% *w/v* BSA, 0.1% *v/v* Triton X-100 for 30 min. The cells were incubated for 2 hours with RNAPII-pSer5 conjugated AlexaFluor488 1:500 (AB196152) at room temperature or RNAPII-pSer2 conjugated AlexaFluor488 1:200 (AB237279) at 4°C overnight. Cells were washed three times with 0.2% *w/v* BSA, 0.1% *v/v* Triton X-100. Cells were washed once with PBS then TBS and fixed for a second time with 4% *w/v* PFA in PBS for 10 min. The cells were washed with PBS three times before being stored in PBS supplemented with 0.02% *w/v* sodium azide. The coverslips were stored at 4°C in the dark. Prior to imaging, the coverslips were washed in PBS twice and placed into an Attofluor cell chamber (Invitrogen, MA, USA). STORM buffer containing 10% *w/v* glucose, 10 mM NaCl, 50 mM Tris-HCl pH 8.0 with GLOX solution consisting of 5.6% *w/v* glucose oxidase and 3.4 mg/mL catalase in 50 mM NaCl, 50 mM Tris-HCL pH8.0 and 0.1% *w/v* β-mercaptoethanol was used for imaging. STORM images were obtained using a Zeiss Elyra PS.1 system. Sample illumination was achieved using an HR Diode 488-200 (100mW) laser with 7-12kW/cm^2^ power density exposed to the sample. The built-in multi-band dichroic mirror MBS 405/488/642 (Carl Zeiss 1784-996) was used to reflect the excitation laser beams onto the sample. Imaging was performed using a 100x NA 1.46 oil immersion objective (Carl Zeiss alpha Plan-Apochromat, 420792-9800-00) and to reduce background fluorescence levels highly inclined and laminated (HILO) illumination was used. To capture fluorescence signal, a BP 420-480/BP 495-550/LP 650 multi-bandpass emission filter (Carl Zeiss 1769-207) was used. The final image acquisition was achieved using an Andor iXon Ultra 897 EMCCD camera with a 30 ms exposure for 20000 frames. Zeiss Zen Black software was used to detect blinking events with a 9-pixel mask and a signal-to-noise ratio of 6. Final molecule positions were determined through fitting a 2D Gaussian function. Model-based cross-correlation drift correction was used on the STORM image. For AlexaFluor488 the mean value of localisation procession was 30 nm. Further analysis was conducted using ClusDoC (49) in MatLab using the molecule localisation table generated from Zeiss Zen black software. A minimum of 30 cells were imaged and processed for ClusDoC analysis.

ClusDoC

STORM data was further analysed using ClusDoC (https://github.com/PRNicovich/ClusDoC) (49). The region of interest (ROI) was selected for the nucleus for cluster analysis. To identify the r max the Ripley K function was calculated for the ROI and this was used for DBSCAN analysis. Minimum cluster size was set to 5 molecules with a smoothing value set at 7 and the epsilon value set at the mean localisation precision value for the dye. Default values were used for the remaining parameters.

### X-Ray Irradiation

CHO and HEK293 cultures were irradiated with 6 Gy dosage using a CIX1 (Xstrahl, Camberly, UK) with the settings; 160 kV, 6.250 mA, 2 rotations/min. The cultures were sampled at 10 min, 30 min and 4 h for alkaline comet assays. n = 3 biological replicates were irradiated and investigated.

### Tryptic Peptide Mapping Using Ultra-Performance Reverse Phase Coupled to ESI Tandem Mass Spectrometry (Peptide Mapping-LC-MS/MS)

All samples were diluted to 5 mg/mL using HPLC grade water. The diluted sample was mixed with a denaturing mix of 7.2M guanidine HCl, 90 mM tris, 0.1 mM EDTA, and 30 mM DTT and incubated at 37°C for 30 min for denaturation and reduction. The alkylation was performed by adding 500 mM iodoacetamide, followed by incubation at ambient temperature for 30 min in the dark. Alkylated samples were then buffer exchanged into 6 M urea using a micro-dialysis device. Samples were diluted in digestion buffer, and freshly reconstituted Trypsin Gold (Promega, WI, USA) was then added to the protein solution at a 1:10 ratio (w/w protease /protein) and incubated at 37 °C for 3.5 h. Proteolysis was stopped by adding 10% trifluoroacetic acid (TFA). The resulting peptide mixtures were then separated by LC-MS/MS using a Waters Acquity UPLC system equipped with a Waters Acquity BEH C18 column, mobile phase A (0.02% TFA in HPLC water), mobile phase B (0.02% TFA in acetonitrile). Mass measurements were performed on an Orbitrap QExactive HF-X mass spectrometry (Thermo Fisher Scientific, MA, USA) mass spectrometer using a data-dependent acquisition method and higher energy collisional dissociation fragmentation. Data were analysed using PMI-Byos v5.9 software (Protein Metrics, CA, USA).

### Microchip Capillary Electrophoresis (MCE) via GXII Touch

A high-throughput analysis of mAbs was performed on the LabChip GXII Touch instrument (Perkin Elmer) using an established protocol (50). Sample preparation procedure was adapted from the procedure outlined by Chen *et al*. (51). An aliquot of 5 µL of sample at 1 mg/mL was mixed with 35 µL of non-reducing sample buffer in a 96 well PCR plate. The non-reducing sample was prepared by mixing 700 µL HT protein express sample buffer with 24.5 µL of 250 mM iodoacetamide (IAM) solution. Samples were then incubated at 70°C for 15 mins, after which 70 µL of deionised water was added to each sample before loading on to the instrument for analysis.

### Statistical Analysis

Statistical analyses were performed using GraphPad Prism (GraphPad, USA), v9.4.0 or R v4.2.3. Results were expressed as mean values. The Kruskal-Wallis test was used for comparisons between 3 groups or more with either Dunn’s multiple comparison tests or Benjamini, Kreiger and Yekutieli method for adjustments or one-way ANOVA with Turkey’s multiple comparison test. For comparisons between 2 groups the Mann-Whitney U test was used. Benjamini and Hochberg’s method was used to generate the false discovery rate (FDR) for transcriptomic analysis. Only two-sided p-values were calculated. Unless otherwise stated, experiments were performed using two independent biological replicates per condition for cell line A and cell line B except one biological replicate for cell line B high VCD. X-ray irradiation experiments and comparisons of freshly cultured versus thawed cells were conducted using three biological replicates except for STORM imaging and AFM, which used one biological replicate. The significance levels are indicated by asterisks in each figure between groups: ns p > 0.05; *p < 0.05; **p < 0.01; ***p < 0.001; ****p < 0.0001.

## RESULTS

### High Cell Culture Densities Impact Cell Viability

Two monoclonal antibody-producing CHO cell lines (cell line A and B) were generated and selected, using an established industry-standard cell line development process. Their selection has been made on the base of fed-batch performance in a small scale Ambr15 microbioreactor **(Supplementary Figure S2)**. Both cell lines exhibited robust growth kinetics and product titres consistent with industrially relevant production lines **(Supplementary Figure S2A–D)**. Cell line A displayed an increase of lactate concentration towards the end of the run, not observed in cell line B, suggesting distinct metabolic states **(Supplementary Figure S2E)**. The two cell lines were also characterised by small differences in transgene copy number, heavy and light chain (HC/LC) transcript abundance, and intracellular HC/LC peptide levels **(Supplementary Figure S3A–D)**.

A schematic of the perfusion bioreactor setup is shown in **Figure 1B**. Seven bioreactors were operated for 14 days under two VCD targets: control (90×10^6^ cells/mL) and high (150×10^6^ cells/mL). Cell line A control and high VCD bioreactors displayed comparable cell growth up to day 4 and then diverged to their respective VCD targets until day 14 **(Figure 1C)**. Maintenance of cell densities at the chosen VCD started on day 3 **(Supplemental Figure 4A-B)**. Accordingly, we define a growth phase (days 0–4), during which cultures approached target VCD, and a production phase (days 5–14), corresponding to the most productive stage of the bioreactor run. Viability for cell line A control and high VCD cultures remained >97% until day 7 **(Figure 1C-D)**. After day 7, cell viability declined across all bioreactors and continued to fall, reaching >94% in the control VCD and <91% in the high VCD by the end of the run. This trajectory suggests the downward trend would be predicted to persist beyond day 14. A similar growth kinetic trend was also observed in cell line B **(Supplemental Figure S4C-D).**

The metabolic profiles (glucose and lactate concentrations) of all the bioreactors followed a similar trend **(Supplemental Figure S4E-H)**. Cell line A and cell line B high VCD bioreactors produced higher titres in both the amount of titre retained within the bioreactor (retentate titre) and the amount of titre filtered from the bioreactor (permeate titre) **(Supplemental Figure S5A-D).** The cell specific productivity (qP) was similar between the high and control VCD bioreactors for both cell lines **(Supplemental S5E-H).**

### Transcriptomic Analysis Revealed a Downregulated DNA Damage Response Across the Perfusion Run

To dissect the molecular processing underpinning the loss of cellular viability. Time-resolved transcriptomic profiling was performed to compare cellular processes and pathways between control and high VCD bioreactors across the perfusion run. Sampling spanned both growth and production phases (**Figure 1B)**, enabling analysis of temporal dynamics, cell line identity and VCD target. Principal component analysis (PCA) indicated that samples separated primarily by timepoint along the first principal component (PC1), capturing 37% of the variance, and by cell line along the second principal component (PC2), capturing 19% variance **(Figure 2A)**. Growth phase samples (days 0, 2, and 4) clustered together irrespective of VCD, whereas production phase samples diverged from this cluster. Cell line A and cell line B formed distinct clusters along PC2, indicating inherent cell line differences, but their trajectories over time were highly similar.

**Figure 2.**
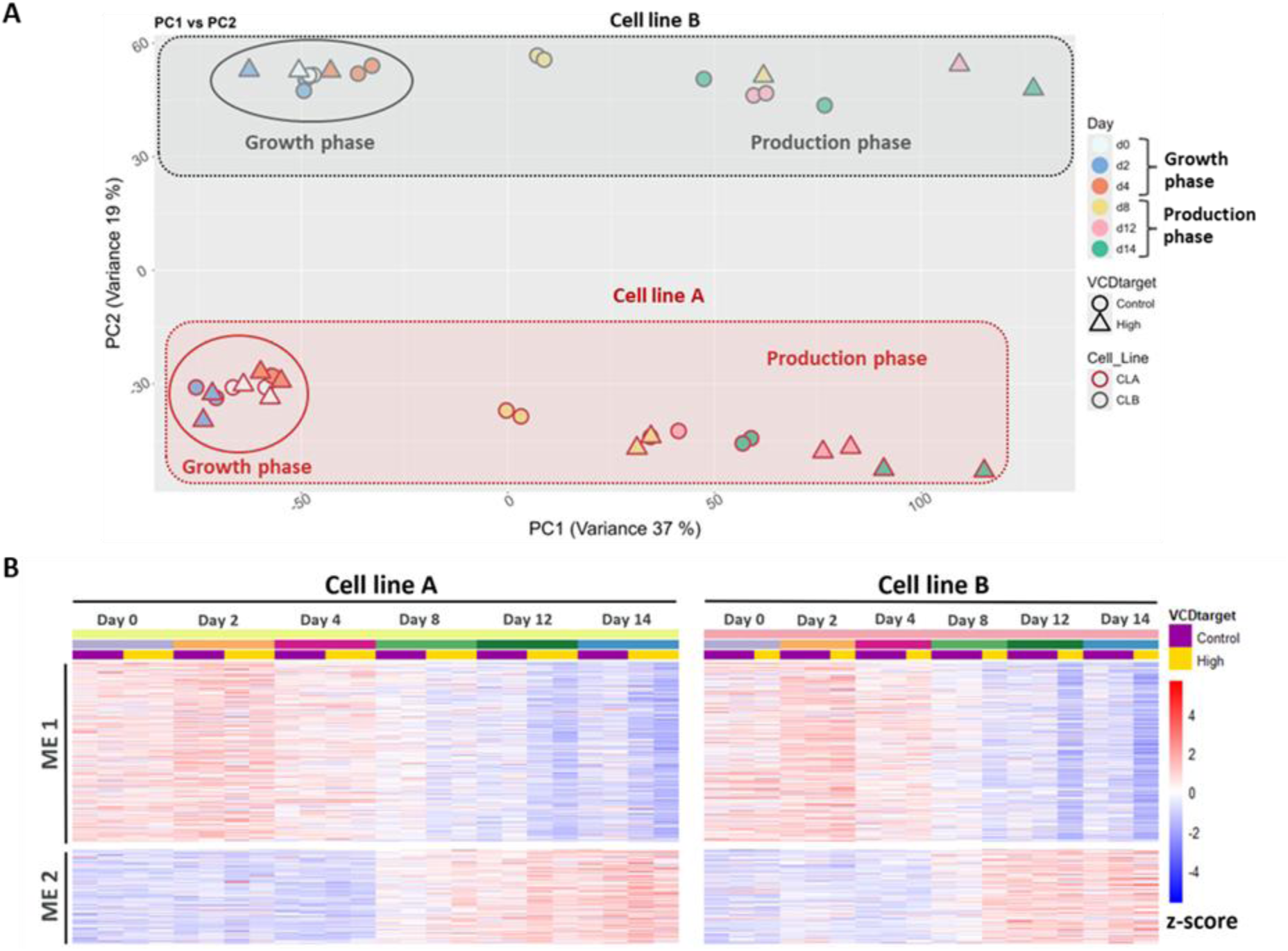
Transcriptomic analysis of CHO cell lines during perfusion bioreactor culture to assess gene expression dynamics under different viable cell density (VCD) conditions. Samples for transcriptomic analysis were collected from each bioreactor on days 0, 2, 4, 8, 12, and 14. (A) Principal component analysis (PCA) of variance-stabilising transformation (VST) normalised counts, with samples represented by day, cell line, and VCD target (high: 150×10⁶ cells/mL, control: 90×10⁶ cells/mL). (B) Weighted gene co-expression network analysis (WGCNA) grouped genes into modules of highly correlated gene expression. Heatmap showing z-score normalised expression of module 1 (ME1) and 2 (ME2) genes in cell line A and B.

The significant resource and time demand for perfusion bioreactor runs limited the number of bioreactors available for this study. To this end, weighted gene co-expression network analysis (WGCNA) (42, 43) was conducted to explore transcriptomic variation across cell line, day and VCD **(Figure 1B)**. A signed network was built from pairwise gene correlations. Then modules were defined by hierarchical clustering with dynamic tree cutting (52), and closely related modules were merged using a stringent threshold derived from the power-law fit (R^2^ = 0.9). This yielded 14 modules in total. After merging highly similar modules with the stringent threshold, 9 final modules were generated **(Supplementary Figure S6A).** Module 1 contained the largest number of genes (3, 497 genes; **Supplementary Figure S6B**) whilst the number of genes in other modules progressively decreased to module 9 (160 genes). Enrichment analysis revealed that module 1 was enriched for Gene Ontology Biological Process (GO:BP) terms related to DNA damage response, gene expression, and DNA repair **(Table 1)**. Module 2 was enriched for processes involved in protein transport, Golgi vesicle transport, and endoplasmic reticulum stress **(Table 2)**. Modules 3, 4, 6, and 7 displayed cell line specific enrichment patterns, featuring processes related to protein localisation, endoplasmic reticulum associated degradation (ERAD) pathway, positive regulation of transcription and proteasomal protein catabolic process **(Supplementary material, Modules_GO_BP)**. Modules 8 and 9 were enriched for terms related to cytoplasmic translation and regulation of gene expression, respectively. Module 5 did not show significant enrichment for any GO:BP terms. Full enrichment analysis for all modules is available in the supplementary material files named Modules_GO_BP.

**Table 1.** Module 1 Gene: Ontology Biological Process (GO:BP) enrichment analysis conducted using enrichR. The top 10 most significantly enriched Gene Ontology Biological Process (GO:BP) terms are shown, ranked by the lowest Benjamini-Hochberg-corrected p-values. Each row lists the GO term, GO:ID, description, and adjusted p-value (Padj). The “Count” column indicates the proportion of genes within the module associated with each ontology. Full enrichment results are provided in the supplementary material (Modules_GO_BP).

| Module 1 Enrichment Analysis |  |  |  |
| --- | --- | --- | --- |
| GO: ID | Description | Padj | Count |
| GO:0006259 | DNA Metabolic Process | 2.93E-45 | 160/288 |
| GO:0000398 | mRNA Splicing, Via Spliceosome | 2.93E-45 | 132/211 |
| GO:0006281 | DNA Repair | 4.12E-44 | 159/291 |
| GO:0000377 | RNA Splicing, Via Transesterification Reactions With Bulged Adenosine As Nucleophile (GO:0000377) | 4.94E-38 | 112/180 |
| GO:0006397 | mRNA Processing | 6.32E-37 | 123/214 |
| GO:0006974 | DNA Damage Response | 6.75E-31 | 168/384 |
| GO:0006396 | RNA Processing | 1.31E-26 | 99/183 |
| GO:0006302 | Double-Strand Break Repair | 1.06E-23 | 90/168 |
| GO:0006261 | DNA-templated DNA Replication | 6.31E-21 | 50/69 |
| GO:0022618 | protein-RNA Complex Assembly | 7.54E-21 | 80/150 |

**Table 2.** Module 2 Gene: Ontology Biological Process (GO:BP) enrichment analysis conducted using enrichR. The top 10 most significantly enriched Gene Ontology Biological Process (GO:BP) terms are shown, ranked by the lowest Benjamini-Hochberg-corrected p-values. Each row lists the GO term, GO:ID, description, and adjusted p-value (Padj). The “Count” column indicates the proportion of genes within the module associated with each ontology. Full enrichment results are provided in the supplementary material (Modules_GO_BP).

| Module 2 Enrichment Analysis |  |  |  |
| --- | --- | --- | --- |
| GO: ID | Description | Padj | Count |
| GO:0006996 | Organelle Organization | 6.09E-09 | 87/418 |
| GO:0010256 | Endomembrane System Organization | 2.83E-08 | 51/196 |
| GO:0015031 | Protein Transport | 2.83E-08 | 69/313 |
| GO:0048193 | Golgi Vesicle Transport | 6.85E-06 | 46/197 |
| GO:0016236 | Macroautophagy | 1.47E-05 | 30/104 |
| GO:0034976 | Response To Endoplasmic Reticulum Stress | 0.000117 | 30/114 |
| GO:0006888 | Endoplasmic Reticulum To Golgi Vesicle-Mediated Transport | 0.000124 | 30/115 |
| GO:0016050 | Vesicle Organization | 0.000199 | 26/94 |
| GO:0016192 | Vesicle-Mediated Transport | 0.000225 | 71/411 |
| GO:0006906 | Vesicle Fusion | 0.000272 | 23/79 |

Pairwise Spearman’s rank correlation coefficient was calculated, using the module eigengene value, to measure distances between the identified modules **(Supplementary Figure S6C)** and hierarchical clustering separated the modules into two broad groups, designated cluster X and cluster Y **(Supplementary Figure S6D).** Cluster X comprised modules 2-5, while cluster Y comprised modules 1 and 6-9. Module gene expression patterns revealed broad trends within each cluster **(Supplementary Figure S6E)**. Cluster X modules displayed low expression until day 4 followed by higher expression post-day 4, whereas cluster Y modules showed the inverse pattern.

None of the modules exhibited significant associations with VCD status, consistent with the comparable growth patterns observed across conditions through day 4 **(Figure 1C; Supplementary Figure S4C)**. Most modules showed clear cell line-specific behaviour, with modules 3–7 displaying significant, cell line-dependent associations, either positively correlated with cell line A and negatively correlated with cell line B, or the inverse **(Supplementary Figure S6F)**. Based on these observations, modules 1 and 2 represented the most numerous and informative gene groups. Their distinct expression profiles, separation into different clusters, and lack of cell line specificity indicated shared and robust expression dynamics across both lines. Therefore, we prioritised these two modules for further characterisation.

Notably, module 1 genes were displayed progressively lower gene expression, reaching a minimum at day 14 **(Figure 2B)**. Whereas module 2 genes showed progressively higher gene expression, peaking at day 14 **(Figure 2B)**. However, genes in module 1 revealed a more pronounced decrease in gene expression in high VCD cultures from day 8 onwards **(Figure 2B).** This observation motivated a day-by-day differential gene expression analysis comparing high and control VCD conditions independently at each time point. To evaluate changes in gene expression associated with different VCD status we used two complementary models: one controlling for cell line variability and one completely omitting the cell line variable **(Supplementary Figure S7A)**. Both approaches produced highly consistent results **(Supplementary Figure S7B).**

Only one differentially expressed gene (DEG) was detected on days 0–4, consistent with the observation that, at these time points, control and high VCD conditions have similar growth profiles **(Figure 1C; Supplementary Figure S4C; Supplementary Figure S7B)**. By contrast, DEGs were detected on days 8, 12, and 14 **(Supplementary Figure S8A–C)**, with day 14 showing the largest shift (100 upregulated, 246 downregulated; **(Supplementary Figure S8C)**. A set of DEGs was shared across the production phase (2 up, 49 down, **Supplementary Figure S9A-B)**. Gene set enrichment analysis (GSEA) using KEGG pathways gave similar results **(Supplementary Figure S10A-C)** at day 8, 12, and 14 for both differential gene expressions models, with enrichment for lysosome, DNA replication, cell cycle, homologous recombination (HR), Fanconi anaemia (FA), and mismatch repair (MMR) pathways in high VCD compared to control VCD bioreactors. Inspecting the gene expression profiles from enriched pathways revealed lysosome associated gene expression increased over time in control and was further elevated at high VCD **(Supplementary Figure S11A-B)**. Conversely, cell cycle, DNA replication, FA, HR, and MMR pathways declined over time and were more markedly downregulated at high VCD **(Supplementary Figure S11A-B)**.

### DNA Damage Accumulates Over Time

Transcriptomic signatures showed a decrease in DNA damage response and signalling transcripts. To explore this further, we quantified phosphorylated histone H2AX (γH2AX), a central marker of double strand breaks, by western blotting and immunofluorescence across bioreactor time points using frozen cell pellets **(Figure 3A–C)**.

**Figure 3.**
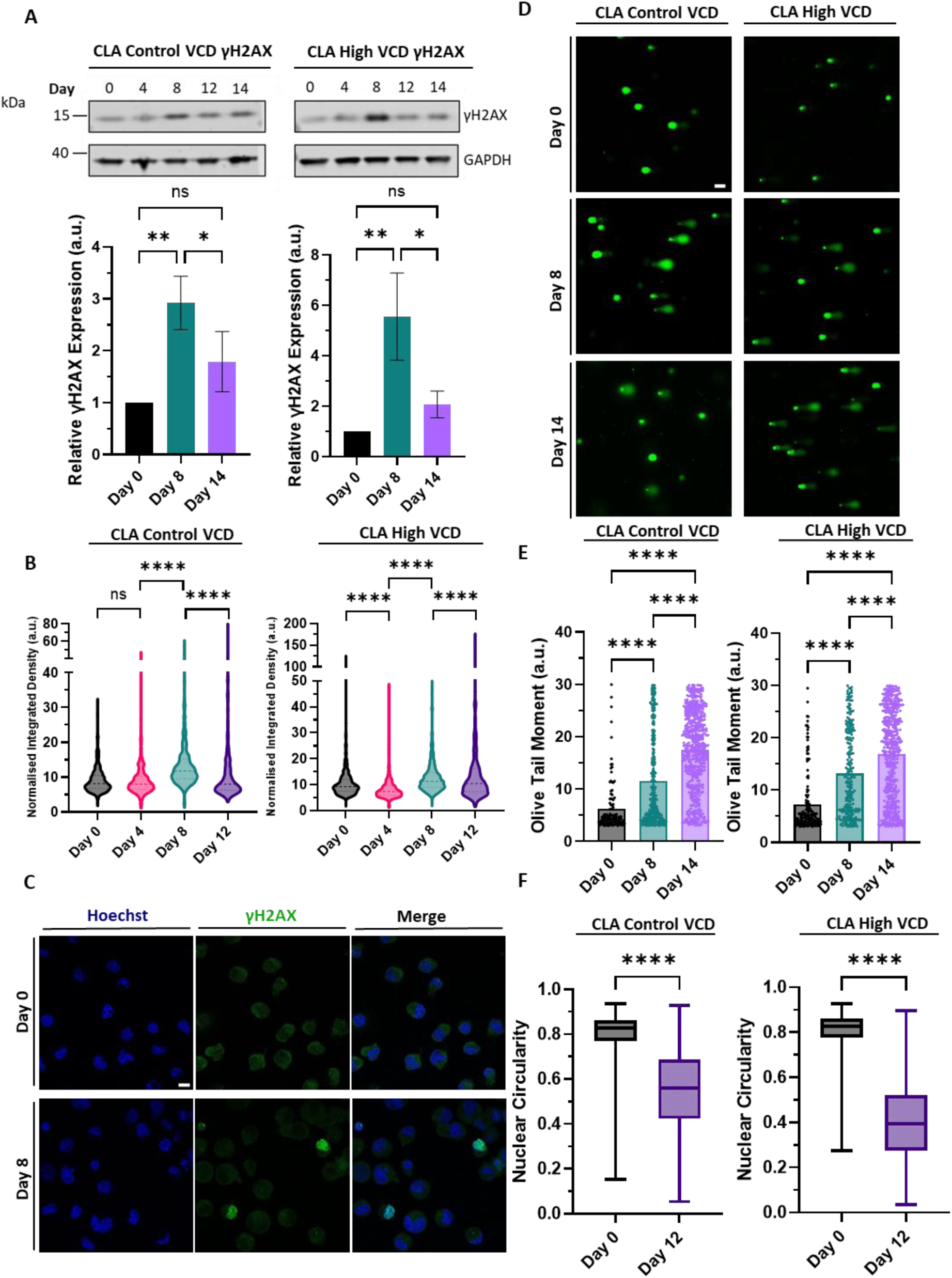
Assessment of DNA damage and nuclear architecture. (A) Representative γH2AX western blots for cell line A (CLA) control and high viable cell density (VCD) bioreactors with corresponding γH2AX normalised to GAPDH densitometry relative to day 0. Error bars show mean ± SD, n = 3 technical replicates, statistics determined by one-way ANOVA with Turkey’s multiple comparisons test. (B) Quantification of γH2AX integrated density normalised to background fluorescence. CLA control VCD: n = 1061/1055/1024/1045, CLA high VCD: n = 1113/1130/1124/1056, statistics determined by Kruskal-Wallis test with Dunn’s multiple comparisons test. (C) Representative confocal imaging of CLA bioreactor derived cells stained against γH2AX (green), with DNA staining shown in blue. Scale bar = 10 µm. (D) Representative alkaline comet assay images, DNA stained with Vista green dye. Scale bar = 20 µm. (E) Alkaline comet assay analysis measuring olive tail moments in CLA control and high VCD bioreactors. Data thresholding was used to remove undamaged and non-viable cells. Full data available in Supplementary Figure S10C. CLA control VCD: n = 121/293/488, CLA high VCD: n = 158/238/471, statistics determined by Kruskal-Wallis test with BKY multiple comparisons test. (F) Quantification of nuclear circularity, CLA control VCD: n = 3214/3124, CLA high VCD: n = 3199/3206, statistics determined by Mann-Whitney U test. In all cases, two biological replicates were performed per condition. Significance definitions, ns p >0.05, *p < 0.05, ***p < 0.001 ****p < 0.0001.

Due to logistical constraints, cellular physiology assays were conducted on frozen bioreactor aliquots. To evaluate potential thawing effects, we compared freshly harvested cells with cells thawed immediately after freezing. No substantial differences were detected in immunofluorescence staining or nuclear circularity between the two groups **(Supplementary Figure S12A-D).**

In cell line A, γH2AX intensity peaked at day 8 and declined thereafter in both western blotting and immunofluorescence imaging **(Figure 3A–C)**, implying either DNA repair or diminished DNA damage signalling. In western blot densitometry and immunofluorescence imaging analysis, this trend was statistically significant in both control and high VCD conditions. Cell line B exhibited similar trends **(Supplementary Figure S13A-B).** To investigate this further, we assessed DNA damage accumulation and repair capacity using the alkaline comet assay, which is particularly sensitive to strand breaks and alkali-labile sites (52). These alkali-labile sites represent key DNA lesions that are specifically revealed under alkaline assay conditions (53).

Alkaline comet assays on days 0, 8, and 14 revealed increasing olive tail moments **(Figure 3D–E)**, indicating time dependent DNA damage. In cell line A control VCD, olive tail moments values rose from 6.2 (day 0) to 11.54 (day 8) and 17.45 (day 14); high VCD showed 7.2, 13.18, and 16.89, respectively **(Figure 3D–E)**. Cell line B exhibited similar trends **(Supplementary Figure S13D–E)**. Collectively, these findings point to substantial accumulation of unrepaired DNA damage by day 14, attributable to a downregulated DNA damage response.

Unrepaired DNA damage has been shown to disrupt cellular biomechanics, particularly nuclear integrity (54–56) which impacts cellular viability. To assess this, we first quantified nuclear circularity. In cell line A at control VCD, circularity declined from 0.80 (day 0) to 0.55 (day 12); at high VCD, it fell from 0.80 to 0.40 **(Figure 3F)**, indicating altered nuclear organisation. We next measured nuclear stiffness via AFM mechanical mapping. Although the DNA damage response is typically associated with decreased nuclear stiffness, control VCD showed a modest, non-significant increase (10.17 kPa to 11.20 kPa), whereas high VCD increased significantly from 11.87 kPa to 16.80 kPa by day 12 **(Figure 4A–B)**, paradoxically supporting a disrupted DDR phenotype.

**Figure 4.**
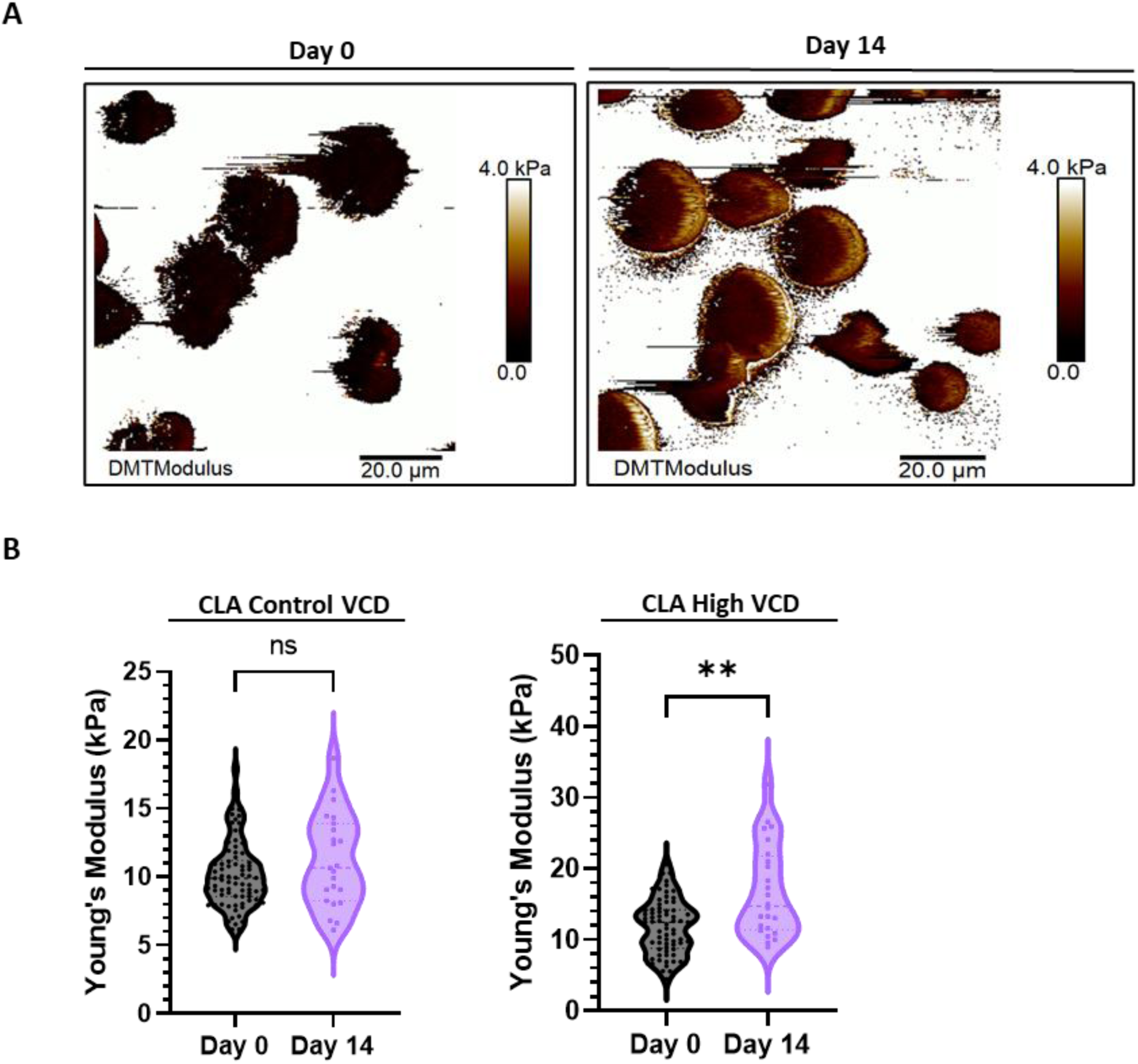
AFM measurements of cell line A control and high viable cell density (VCD) bioreactor derived cells. (A) Representative atomic force microscopy (AFM) nanomechanical map of cells at day 0 and day 14. Scale bar = 20 μm. (B) Young’s moduli values for cell line A (CLA) control and high VCD over time. CLA control VCD: n = 81/23, CLA high VCD: n = 69/24, statistics determined by Mann-Whitney U test. One biological replicate was performed per condition. Significance definitions, ns p >0.05, **p < 0.01.

### Transcriptional Reprogramming is Triggered During the Perfusion Bioreactor Process

DNA damage is closely coupled to RNA polymerase II (RNAPII) regulation and cellular transcription (57, 58). We quantified the levels of RNAPII-pSer5 (initiation) and RNAPII-pSer2 (elongation) proteins by western blotting and immunofluorescence. A significant decrease in RNAPII-pSer5 and RNAPII-pSer2 levels was observed over time in control and high VCD bioreactors **(Figure 5A-F)**. Moreover, total Rpb1 (main RNAPII subunit) protein also decreased **(Figure 5A)**, indicating a global reduction in RNAPII not limited to phosphorylation state. Notably, these changes did not affect titre **(Supplementary Figure S5A–D)** or mAb HC/LC transcript levels **(Supplementary Figure S14A–G)**.

**Figure 5.**
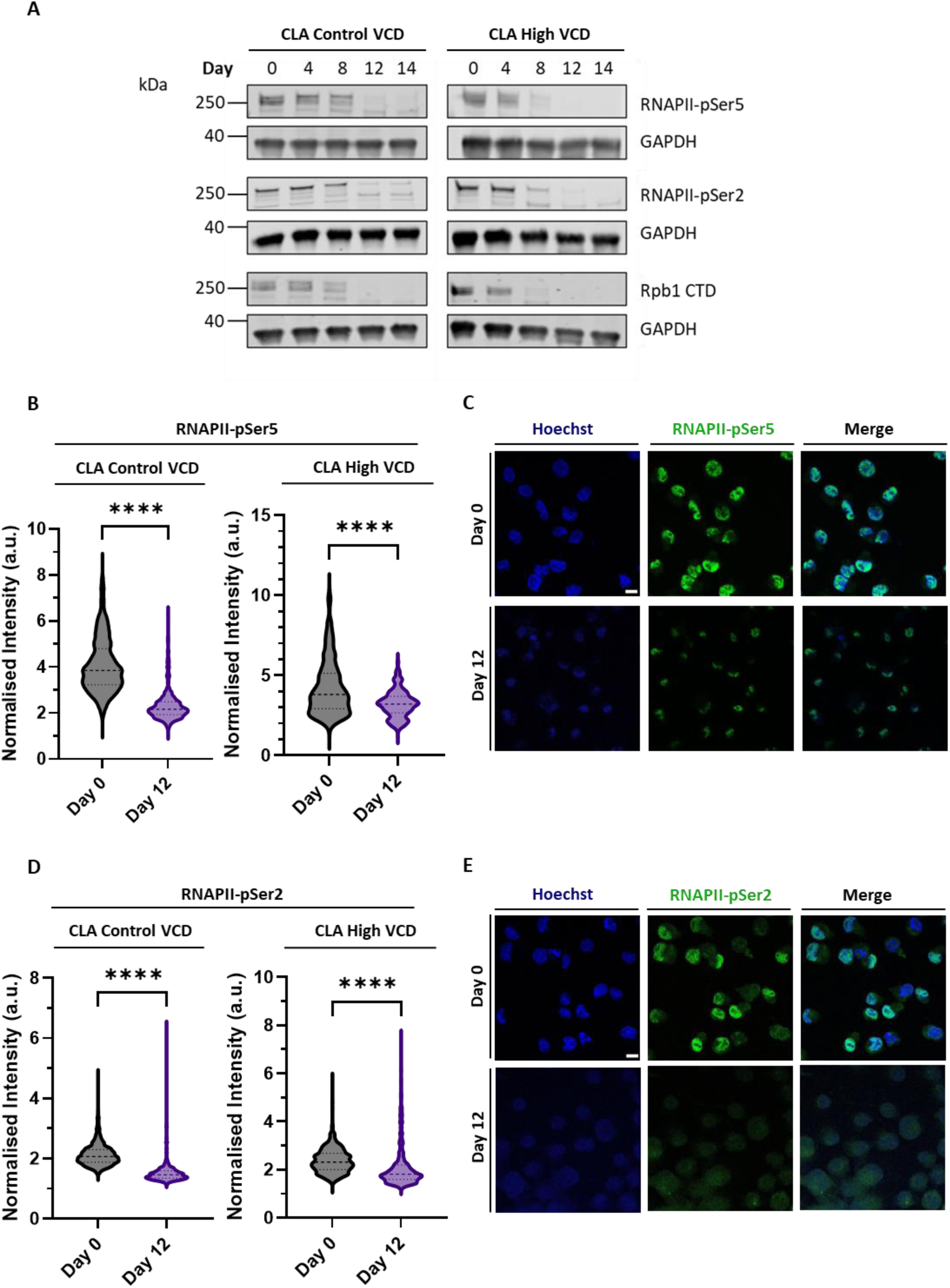
RNAPII analysis by western blot and immunofluorescence. (A) Representative RNA polymerase II (RNAPII)-pSer5, RNAPII-pSer2 and Rpb1 C-terminal domain (CTD) western blots for cell line A (CLA) control and high viable cell density (VCD) bioreactors. (B) Quantification of RNAPII-pSer5 fluorescence intensity normalised to background fluorescence. CLA control VCD: n = 1091/1057, CLA High VCD: n = 523/521, statistics determined by Mann-Whitney U test. (C) Representative confocal imaging of bioreactor derived cells stained against RNAPII-pSer5 (green), with DNA staining shown in blue. Scale bar = 10 µm. (D) Quantification of RNAPII-pSer2 fluorescence intensity normalised to background fluorescence. CLA control VCD: n = 1062/1022, CLA High VCD: n = 992/1101, statistics determined by Mann-Whitney U test. (E) Representative confocal imaging of bioreactor derived cells stained against RNAPII-pSer2 (green), with DNA staining shown in blue. Scale bar = 10 µm. Two biological replicates were performed per condition. Significance definitions, ****p < 0.0001.

To explore this phenotype at a single molecule resolution, we applied STORM, a super-resolution microscopy technique, to quantify RNAPII-pSer5 and visualise transcription hubs via molecular clustering (47), a readout of nuclear transcriptional organisation and cellular state (48, 59). RNAPII-pSer5 molecules declined from day 4 to day 12 in cell line A control VCD (5, 884 to 3, 176) and more sharply in high VCD (16, 821 to 2, 874) **(Figure 6A–B)**, consistent with western blot and immunofluorescence **(Figure 5)**. Spatial analysis using the linearised Ripley’s K function confirmed non-random nuclear clustering (positive deviation; **Supplementary Figure S15A**). Clusters were defined using ClusDoC as ≥5 neighbouring molecules within the mean localisation precision, enabling nuclear cluster mapping **(Figure 6A)**. Cluster numbers decreased from day 4 to day 12 in control (80.32 to 24.66) and more markedly in high VCD (322.7 to 31.71) settings **(Figure 6D)**, whilst mean molecules per cluster slightly decreased and mean cluster area remained stable or decreased modestly **(Figure 6E–F)**. Thus, high VCD cultures exhibited a greater loss of RNAPII-pSer5 clusters, but residual hubs maintained comparable molecular occupancy and size.

**Figure 6.**
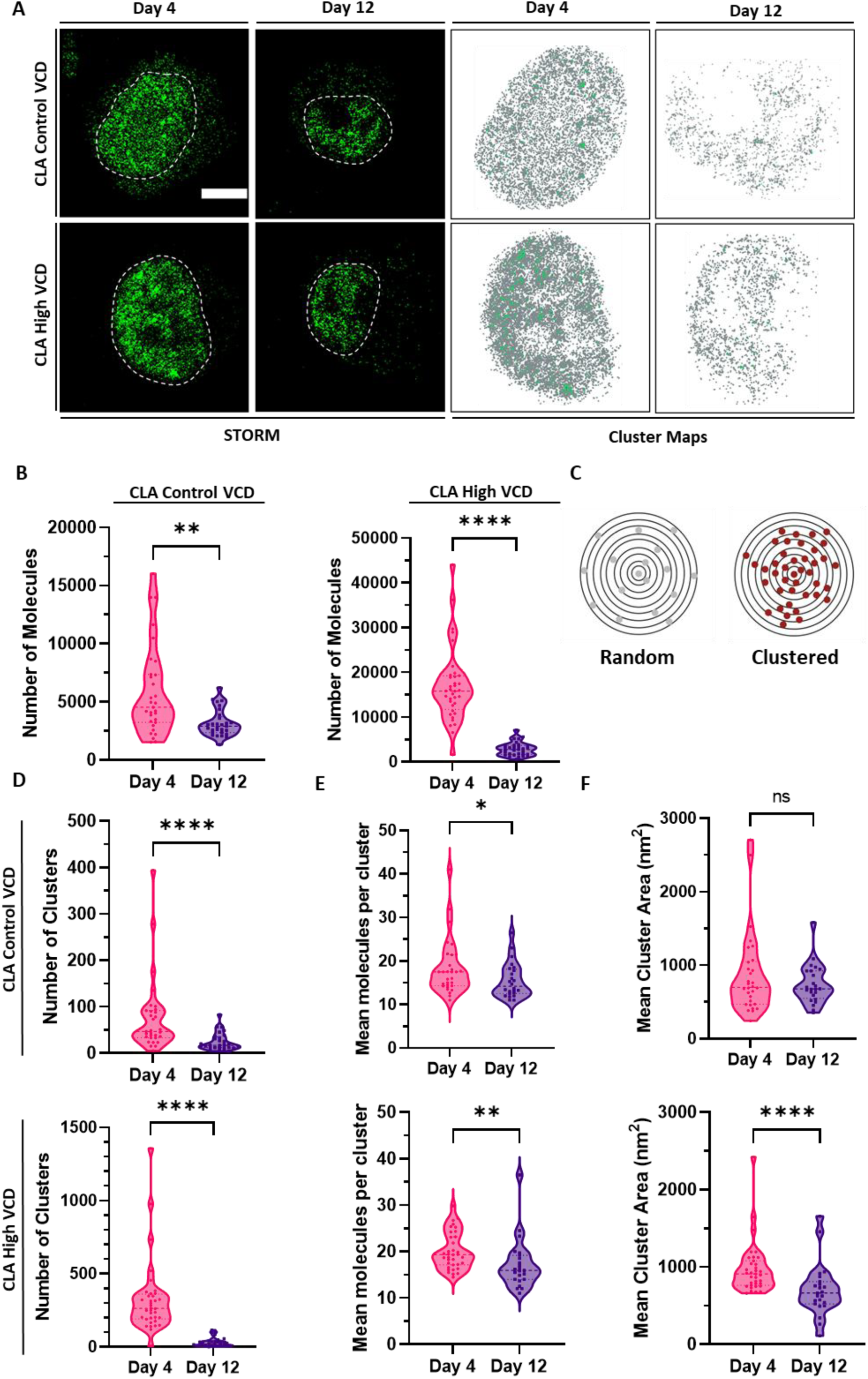
RNA Polymerase II (RNAPII)-pSer5 transcription hub analysis by STORM. (A) Representative STORM image of RNAPII-pSer5 in cell line A (CLA) control and high viable cell density (VCD) conditions at days 4 and 12. The nuclear region (determined by RNAPII-pSer5 fluorescence) was used for ClusDoC analysis (shown as dotted white line). Scale bar = 5 µm. Cluster maps adjacent generated for regions of interests (ROIs), using parameters specified in Materials and Methods. Clustered molecules are shown in green. (B) Total number of molecules detected within ROI, CLA control VCD: n = 31/29 and CLA high VCD: n = 36/31, statistics determined by Mann-Whitney U test. (C) Diagram depicting random distribution and molecular clustering of RNAPII-pSer5. Cluster analysis of CLA control and high VCD RNAPII-pSer5 in the nucleus showing: (D) number of clusters in ROI, (E) mean molecules per cluster and (F) mean cluster area in nm^2^. CLA control VCD: n = 31/29 and CLA high VCD: n = 36/31, statistics determined by Mann-Whitney U test. One biological replicate was performed per condition. Significance definitions, ns p > 0.05, *p < 0.05, **p < 0.01, ****p < 0.0001.

### Extended Perfusion Duration at High VCD Does Not Maintain High Viability

The pronounced accumulation of DNA damage and RNAPII downregulation during perfusion is likely linked to declining culture viability. To probe this further, we conducted an extended perfusion run with cell line A **(Figure 7A-B)**, operating two bioreactors at control VCD and two at high VCD. Here, viability exceeded 90% through day 7, then declined in all vessels **(Figure 7B)**, with high VCD cultures dropping to ∼63% by day 21 and control VCD cultures to 63.5% and 53.8%.

**Figure 7.**
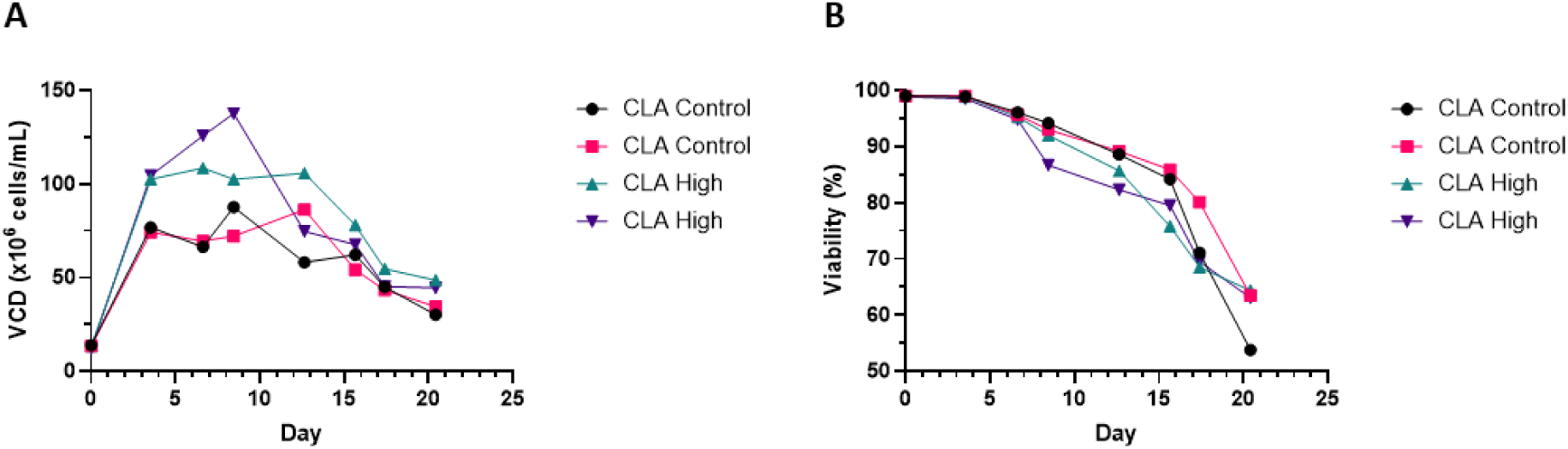
Extended duration perfusion bioreactor culture of cell line A (CLA) to assess long-term growth kinetics and viability. Four bioreactors were operated for 21 days, two at control viable cell density (VCD) target (target: 90×10⁶ cells/mL) and two at high VCD target (target: 150×10⁶ cells/mL), with cultures harvested at day 21. (A) Viable cell density (VCD) profile over time. (B) Cell viability (%) throughout the perfusion culture period. Two biological replicates perfromed per condition.

### CHO Cells Exhibit Attenuated DDR Relative to HEK293 Cells

DNA damage, compounded by mutations in key DDR genes, likely drives reduced viability and suboptimal perfusion performance in CHO cells. We therefore assessed an alternative host, suspension-adapted HEK293 cells to compare acute DNA repair capacity. Here, null-expressing HEK293 and CHO cells were irradiated with 6 Gy, and repair kinetics were quantified by alkaline comet assays over time **(Figure 8A–B)**. At 10 min post-irradiation, olive tail moments were 19.0 (CHO) and 13.0 (HEK293); by 30 min, CHO increased to 25.8 while HEK293 decreased to 5.6. At 4 h, CHO declined modestly to 24.4, whereas HEK293 fell further to 2.3. Controls were stable across timepoints. These DNA repair kinetics indicate an intact DDR in HEK293 and an attenuated DDR in CHO.

**Figure 8.**
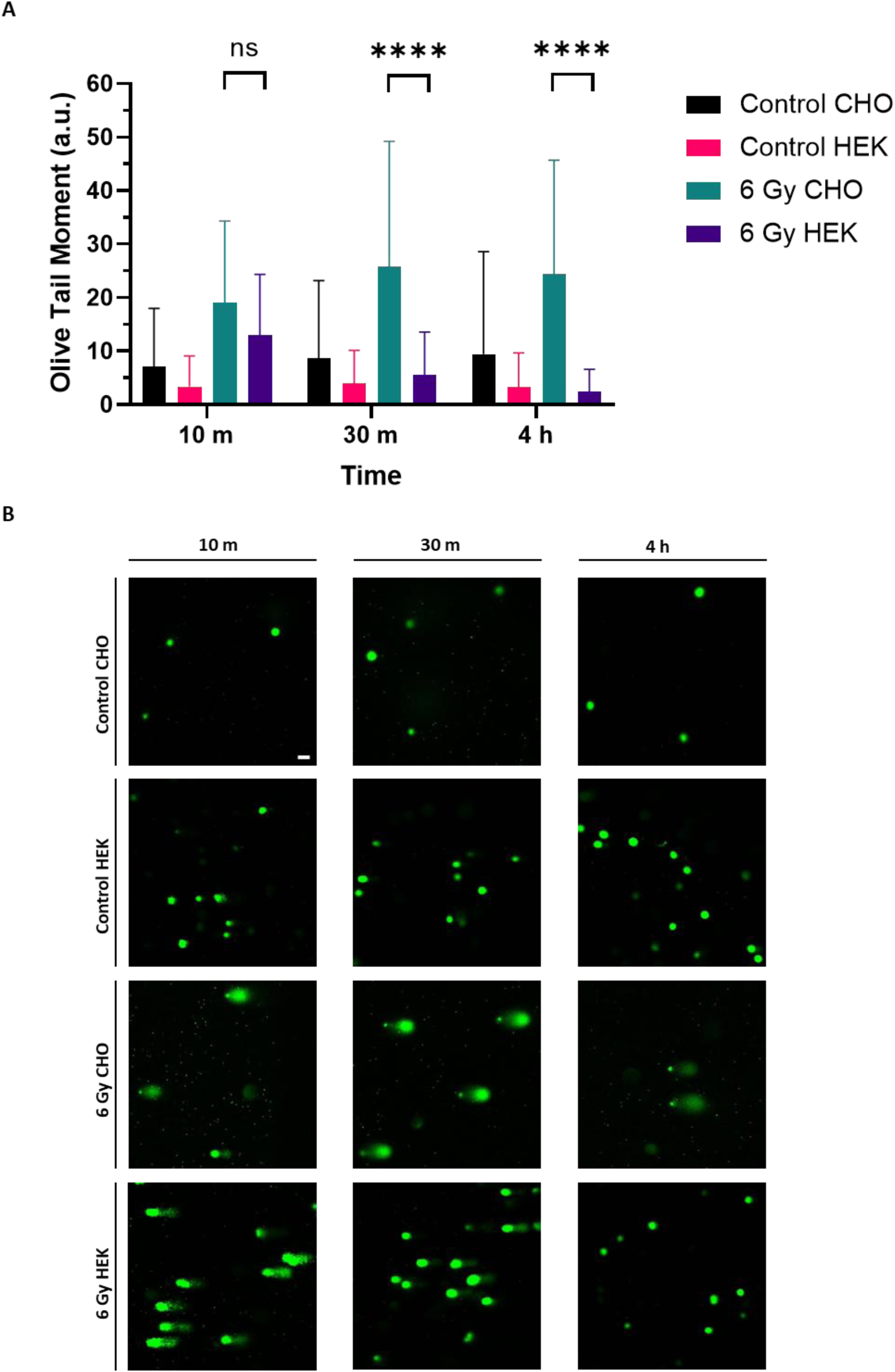
CHO and HEK293 cell DNA repair kinetics post 6 Grays (Gy) dosage irradiation treatment. (A) DNA damage measured by alkaline comet assay at 10 min, 30 min and 4 h post-irradiation. Control CHO: n = 157/125/156. Control HEK: n = 702/515/824. 6 Gy CHO: n = 173/135/133 and 6 Gy HEK n = 377/569/762, error bars indicate mean ± SD, statistics calculated by Kruskal-Wallis test with BKY multiple comparisons test. (B) Representative alkaline comet assay images, DNA stained with Vista green dye. Scale bar = 20 µm. Three biological replicates were performed per condition. Significance definitions, ns p > 0.05 and ****p < 0.0001.

## DISCUSSION

The increasing demand for biopharmaceuticals and the design of more complex molecules for clinical use has challenged previous manufacturing processing paradigms (60–63). Perfusion bioreactors help support these demands by delivering improved cellular growth, productivity and superior product quality compared to conventional fed batch approaches (8, 60). Despite these advantages, extending production phases during perfusion culture remains hampered by declining viability at both moderate (14, 18–20) and high cell densities (17, 21, 64). This decline in viability has been attributed to process related issues (9–11) and cell stressors (3, 12). Data presented here supports a model in which CHO cells experiencing cumulative DNA damage progressively suppress DNA repair and DNA damage signalling pathways rather than activating a sustained DDR, leading to transcriptional collapse and ultimately loss of viability.

The DDR is critical for regulating cellular health and survival (65). Transcriptomic data described here revealed a broad downregulation of genes involved in the DDR during the production phase of the perfusion process. Consistent with this observation, previous reports show that DNA repair was downregulated in the declining phase of batch culture processes, which was proposed to increase genomic instability (66, 67). To further probe DNA damage in perfusion culture, the level of γH2AX protein, a canonical marker of DNA double-strand breaks was measured (68). Here we observed a transient increase followed by a decline in γH2AX levels, an observation that suggests DNA damage was detected followed by either DNA lesion repair or diminished DNA damage signalling **(Figure 3A-C).** Alkaline comet assays revealed persistent DNA lesions despite the apparent loss of the γH2AX signal **(Figure 3D-E)**. This discordance suggests that DDR signalling becomes uncoupled from effective DNA repair, permitting the accumulation of unresolved DNA damage despite the absence of sustained γH2AX marking. In addition, a reduction in proliferation arising from the downregulation of cell cycle and DNA replication genes could contribute to the reduced DDR expression **(Supplementary Figure S11A-B).** However, the perfusion culture continues to proliferate to reach the intended VCD targets after the bioreactor bleed and given the substantial DNA damage observed from the alkaline comet assays, this explanation appears unlikely. Instead, the data suggests impaired DNA repair capacity and disrupted DDR signalling.

Biophysical measurements described in this study provide further evidence for impaired DDR in perfusion-derived CHO cells **(Figure 4)**. In contrast to the chromatin decondensation and nuclear softening that typically accompany DDR activation (56, 69), perfusion-derived cells exhibited increased nuclear stiffness. This is notable given that bioreactor cultures experience shear stress, which can induce DNA fragmentation via mechanical forces (70, 71). Nuclear stiffening under perfusion conditions may reflect two non-mutually exclusive mechanisms: elevated stiffness may exacerbate damage accrual by abrogating the protective, stress-induced softening response normally elicited by mechanical perturbation (69), or the stiffening may represent adaptive remodelling to bioreactor-associated shear stress (72) consistent with the more pronounced increase in nuclear stiffness observed at higher viable cell densities. Therefore, cells with stiffer nuclei survive through bioreactor induced selection.

DNA damage can result from several perfusion-associated stressors, including oxidative stress (73), hypoxia (74) and shear stress (11). Oxidative stress arises from excess generation of reactive oxygen species (ROS) (75, 76) which directly damages DNA inducing single and double-strand breaks, DNA-protein crosslinks, and base modifications (77). Hypoxia can further elevate ROS production (78) and downregulate DNA repair pathways (74). Shear stress can induce DNA fragmentation via mechanical forces (79, 80). Given that these stressors are known to induce genotoxic effects it is perhaps surprising that we observed a marked downregulation of the DDR. Based on these findings, we propose that attenuating the DDR may confer a short-term survival advantage to cells in the bioreactor. This advantage likely arises from the high metabolic and energetic demands of sustained proliferation combined with strong promoter-driven transgene expression. This is because the DDR is energetically costly (3, 81) and restrains cell growth through DNA damage-induced cell cycle arrest (82, 83). Suppressing the response may free critical energy resources. However, this benefit is transient; unrepaired DNA lesions accumulate, genomic instability escalates, and cultures ultimately fail to maintain high viability, thereby limiting the duration of the perfusion production phase.

Beyond nuclear mechanics, the relationship between DNA damage and transcription provided further mechanistic insight into the observed viability decline. Transcription and the DDR are interlinked processes under stringent regulatory control (84). DNA damage induces transcriptional repression to prevent RNAPII collision with DNA repair machinery and transcription of damaged DNA sequences. This repression occurs through RNAPII stalling at sites of DNA damage, where RNAPII senses a steric hindrance caused by DNA lesions (57, 85–88). The stalling stimulates ubiquitination of RNAPII (84) and upon DNA repair failure, ubiquitinated RNAPII is degraded by the proteasome (89).

To explore this, RNAPII levels were analysed by western blot and immunofluorescence **(Figure 5)**, which revealed progressive downregulation of total and phosphorylated versions of RNAPII over the course of the perfusion culture. STORM analysis supported this observation revealing a substantial reduction in transcription hubs as measured by RNAPII-pSer5 clustering. Interestingly, the remaining hubs preserved comparable cluster area and RNAPII-pSer5 molecule counts **(Figure 6)**, indicating that surviving transcription hubs maintain their structural integrity despite the overall RNAPII downregulation. Regardless of the widespread loss of transcription hubs, the maintenance of fewer but functionally intact hubs perhaps suggests expression prioritisation of critical gene expression. This may include the expression of the biotherapeutic chains as HC and LC mRNA levels remained unchanged throughout the perfusion run **(Supplementary Figure 14)**, and neither RNAPII loss nor DNA damage accumulation measurably affected mAb product quality **(Supplementary Tables 2–4)**. These findings indicate that while global transcription is compromised through widespread hub loss, product-critical transcripts potentially remain protected within the surviving transcriptional machinery.

The concurrent DNA damage accumulation and RNAPII downregulation observed in this study predicted that these molecular changes would compromise culture viability during extended perfusion bioreactor runs. Consistent with this framework, the 21-day bioreactor run corroborated a progressive viability decline after day 14 **(Figure 7)**. This decline coincided with decreasing VCD, indicating that the culture could no longer sustain the target cell density as growth deteriorated. Together, these observations support the hypothesis that unresolved DNA lesions progressively impair genomic stability and transcriptional fidelity, ultimately limiting perfusion culture longevity.

CHO-K1 derived cell lines harbour mutations in key DDR regulators (e.g., *ATM* and *TP53*) (25), resulting in basal DDR attenuation consistent with the impaired post-irradiation DNA repair kinetics observed. **(Figure 8)**. This deficit is underscored by comparison to suspension-adapted HEK293 cells (90–93), which exhibit a more robust DDR (94), positioning the DDR as a strategic target for process and host cell engineering. While perfusion-focused efforts have largely prioritised apoptosis modulation via knockout of pro-apoptotic factors (3, 16), these approaches do not address the upstream driver of viability loss, ongoing DNA damage, identified here as a central constraint.

The superior DNA repair performance of HEK293 cells supports their use as a reference framework for DDR enhancement in CHO cells. Consistent with this, Spahn *et al.* showed that correcting select DDR mutations in CHO improves post-irradiation repair and genome stability in extended adherent culture (25). However, complete DDR restoration may be suboptimal for perfusion, as partial attenuation has been proposed to confer genomic plasticity and adaptability under bioreactor stress (3, 25). Therefore, a balanced strategy is warranted, whereby DDR capacity is elevated sufficiently to limit DNA damage-driven viability decline while preserving adaptive potential, for example through targeted reversion of DDR mutations guided by HEK293 orthologs or tuneable modulation of DDR gene expression through the use of controlled overexpression or inducible switches. This would enable a calibrated enhancement of DNA repair proficiency without eroding advantageous adaptability. Furthermore, DNA damage can be mitigated through optimisation of process conditions, including media formulation and antioxidant incorporation (95–97), alongside numerous other strategies.

Perfusion bioreactor experiments are inherently resource-intensive, constraining the total number of independent long-duration runs that can be performed. Nevertheless, the central observations reported here were highly reproducible across two independent CHO cell lines and two independent perfusion bioreactor campaigns, encompassing experiments conducted across eleven bioreactor vessels. This consistency supports the robustness and generalisability of the identified DNA damage-driven constraints on culture longevity. Moreover, our work highlights the critical need to fully characterise cellular behaviour within these intensified culture environments.

In conclusion, our findings identify progressive DNA damage accumulation, driven by global downregulation of the DDR, as a fundamental bottleneck limiting CHO cell viability in perfusion culture. Across two antibody-producing cell lines operated at control and high VCD, we show that unrepaired DNA lesions accumulate alongside increased cellular stiffness, loss of RNAPII, and depletion of transcription hubs; molecular and biophysical hallmarks that precede and predict viability decline. An extended perfusion run confirms that this damage-driven decline continues beyond day 14, establishing DNA repair capacity as a key determinant of perfusion longevity. Together, these data position perfusion bioreactors as a high-stress regime that exposes the limits of CHO genomic plasticity. The markedly more efficient DNA repair observed in HEK293 cells further underscores the intrinsically attenuated DDR in CHO cells. Collectively, these findings reframe perfusion culture longevity as a problem of genome maintenance, positioning targeted modulation of DDR pathways as a new axis for CHO host cell optimisation.

## Data Availability

All data are available in the main text or the supplementary materials.

## Supporting information

Modules_GO_BP

## Acknowledgements

The authors would like to thank Fabio Zurlo and the Bioprocess Analytics team for titre analysis, Siddique Uddin for protein purification, Christine Ferng and Kerensa Klottrup-Rees for cell line selection discussion, the Cell Line Development team at AstraZeneca and members of the Toseland Lab at the University of Sheffield for their support and valuable discussion. We wish to acknowledge the Henry Royce Institute for Advanced Materials. OpenAI GPT-5 was used for structuring the text under the supervision of the authors.

## Author Contributions

S.N.S., R.K.M., and C.P.T., conceived and conceptualised the project. N.B.H., S.N.S., R.K.M., and C.P.T., designed experiments. N.B.H., C.S.F., R.K.M. performed cell line generation. M.A., and S.D., performed perfusion bioreactor runs and sample collection. D.R., performed product quality analysis. R.E., performed RNA extraction and library preparation. L.G., and N.B.H performed transcriptomic and bioinformatic analysis. N.B.H., performed cell line characterisation experiments, western blot, immunofluorescence, comet assay and STORM data acquisition/analysis, AFM analysis and irradiation experiments. L.W., provided STORM microscope and guidance. C.P.T., performed AFM. C.P.T., R.K.M., S.N.S., D.A.B., I.M.S., and D.H., acquired funding. N.B.H., C.P.T., R.K.M., and S.N.S., wrote the manuscript. D.H., and K.L., supported the project. D.H., L.G., D.A.B., I.M.S., S.N.S., R.K.M., and C.P.T., reviewed the manuscript.

## CRediT authorship contribution statement

**Rajesh K Mistry:** Conceptualization, Methodology, Supervision, Funding acquisition, Writing – original draft, Writing – review & editing. **Si Nga Sou:** Conceptualization, Supervision, Funding acquisition, Writing – original draft, Writing – review & editing. **Christopher P Toseland:** Conceptualization, Methodology, Supervision, Funding acquisition, Writing – original draft, Writing – review & editing. **Noah B Hitchcock:** Data curation, Formal analysis, Methodology, Visualisation, Writing – original draft. **Maxine Annoh:** Methodology. **Samik Das:** Methodology. **Cristina Sayago Ferreira:** Methodology. **Daniel Ray:** Methodology. **Ramy Elgendy:** Methodology. **Lin Wang:** Resources. **Ken Lee**: Resources. **Luigi Grassi:** Data curation, Writing – review & editing. **Ian M Sudbery:** Data curation, Writing – review & editing, Funding acquisition. **Dan A Bose:** Writing – review & editing, Funding acquisition. **Diane Hatton:** Writing – review & editing, Funding acquisition.

