## Supplementary figures for "DNA Damage Driven Viability Loss and Transcriptional Reprogramming in Chinese Hamster Ovary Cell Perfusion Culture"

**Supplementary Figure S1**

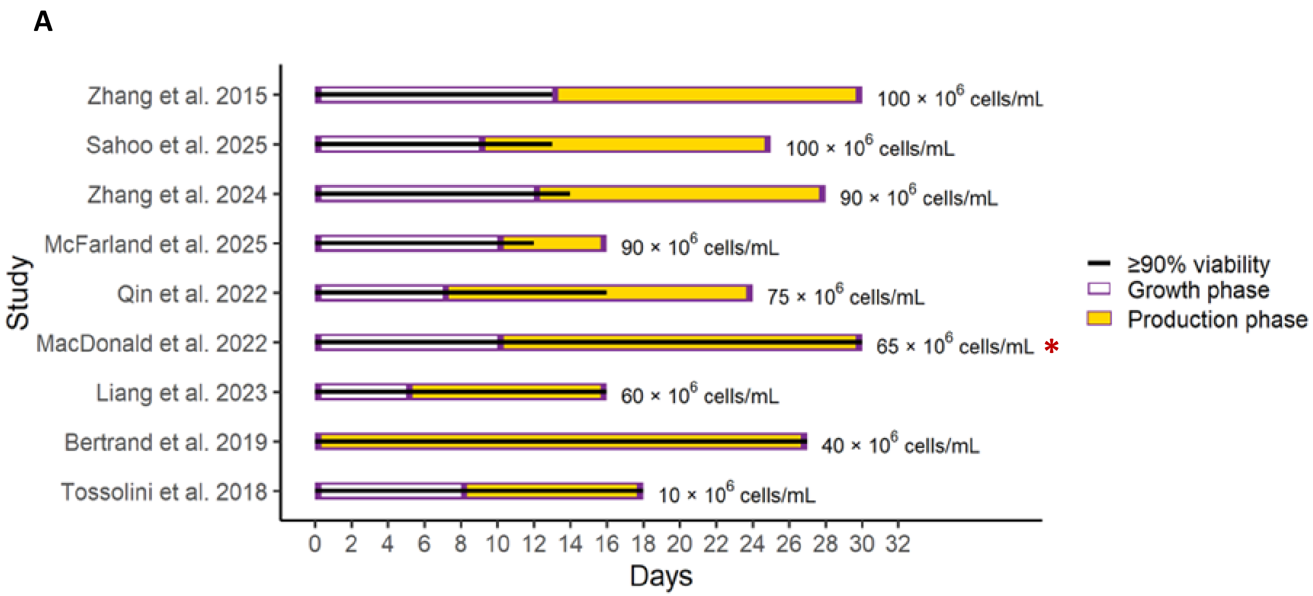


**Supplementary Figure S1. A comparison of published perfusion culture durations.** Each horizontal bar is a representative perfusion bioreactor run study. Purple outlined segments depict the growth phase, the yellow segments indicate the duration of the production phase, and the black line marks the period with ≥90% cell viability. The approximate VCD target for each study is labelled adjacent to each bar. The asterisk highlights the study where an engineered pro-apoptotic gene knockout cell line was investigated.

**Supplementary Figure S2**

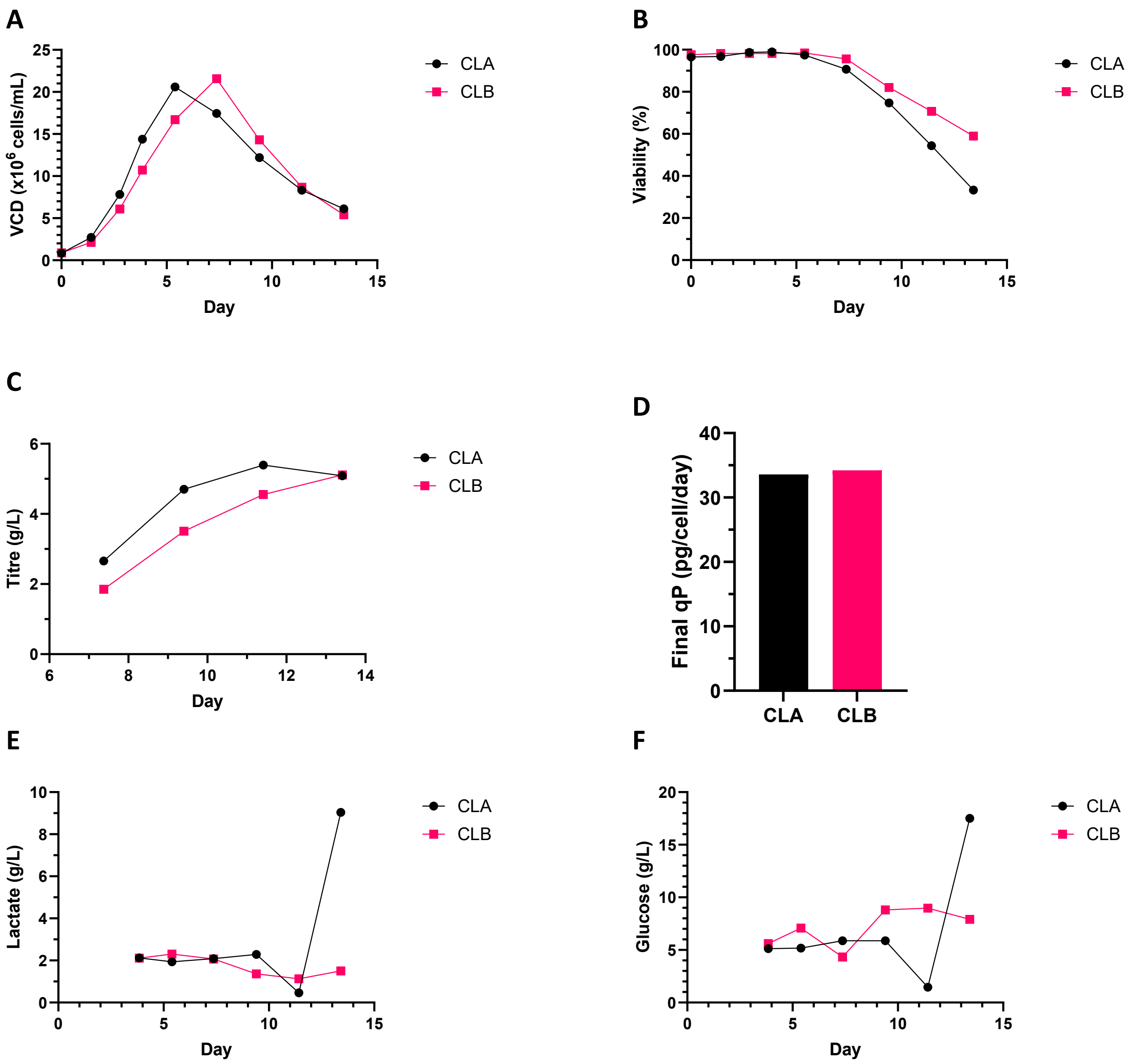


**Supplementary Figure S2. Fed-batch clonal screening of Cell line A and Cell line B in an Ambr15 (automated micro-bioreactor) system.** Clones were cultured in fed-batch mode for 14 days to assess growth, viability, productivity, and metabolic profiles. (A) Viable cell density (VCD) profile over time. (B) Cell viability (%) throughout the culture period. (C) Product titre (g/L) over time collected on days 7, 9, 11 and 14. (D) Final specific cell productivity (qP,) at day 14. (E) Lactate concentration in culture supernatant. (F) Glucose concentration in culture supernatant. One biological replicate performed per condition.

#### Supplementary Figure S3

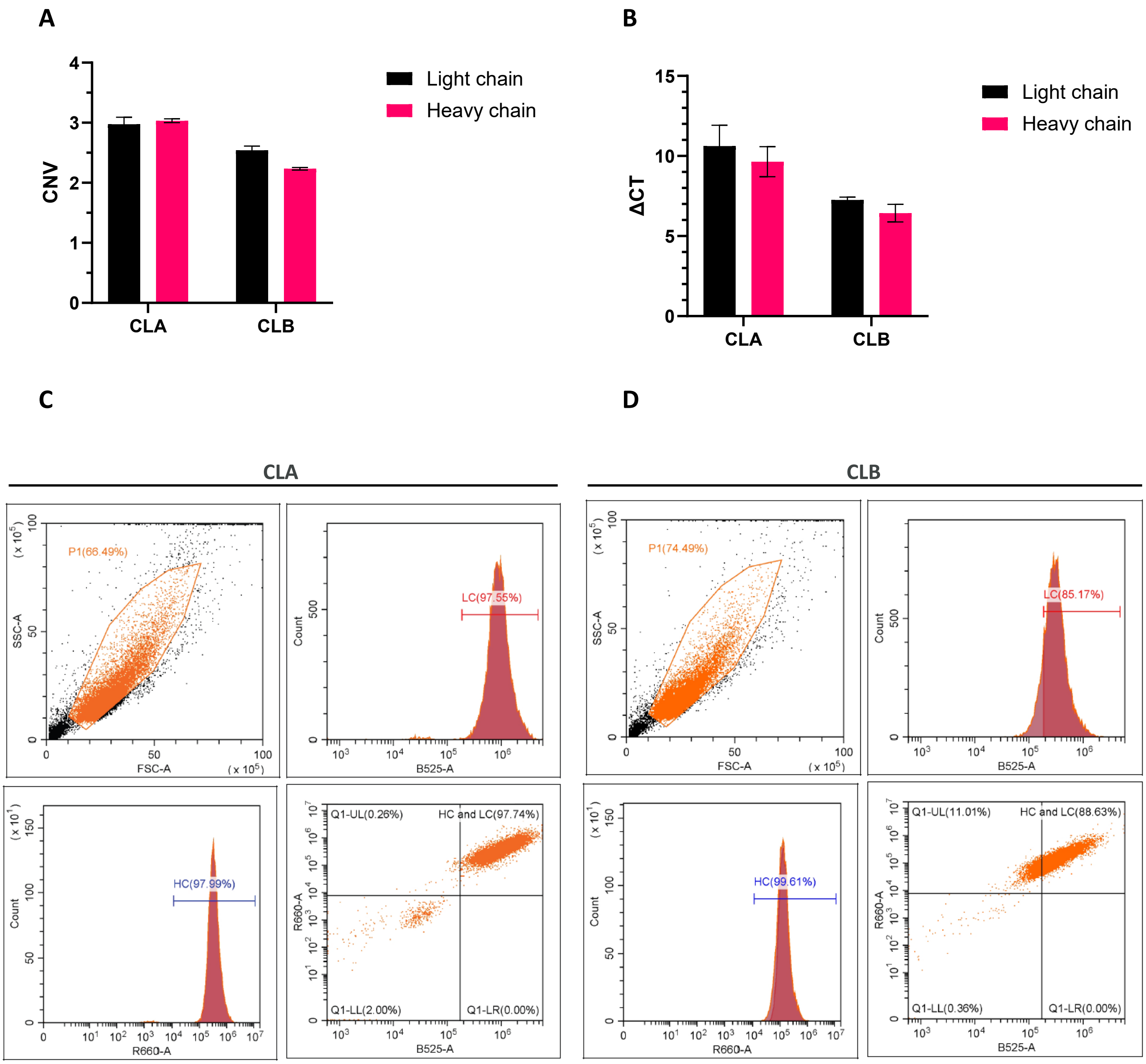


**Supplementary Figure S3. Molecular characterization of Cell line A (CLA) and Cell line B (CLB).** (A) Transgene copy number of heavy (HC) and light chain (LC) genes determined by droplet digital PCR (ddPCR). (B) HC and LC gene expression measured by qPCR and normalised to *Mmadhc* reference gene as described in Materials and Methods. For panels (A) and (B), data represent mean ± SD of n = 3 technical replicates. (C, D) Intracellular HC and LC protein levels in CLA and CLB, respectively, measured by flow cytometry with intracellular staining. Each panel shows: gating strategy for viable cells (upper left), LC-positive cells (upper right), HC-positive cells (lower left), and dual-positive population analysis (lower right) displaying the proportion of cells positive for both HC and LC. One biological replicate was performed per condition.

#### Supplementary Figure S4

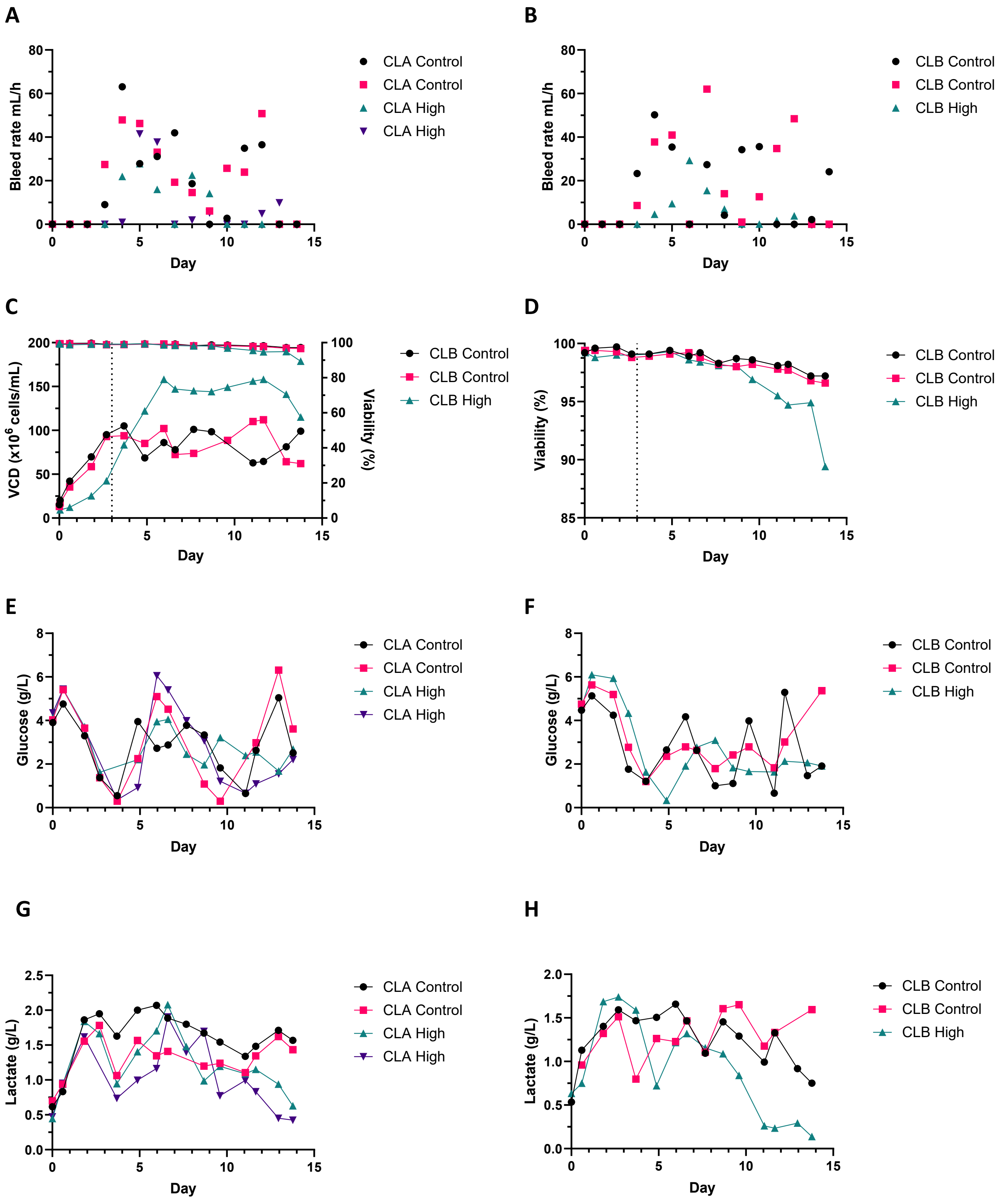

**Supplementary Figure S4. Cell line A (CLA) and Cell line B (CLB) extended perfusion bioreactor run data.** All bioreactors were operated with bioreactor bleed initiation on day 3 (indicated by dashed line. Control viable cell density (VCD) target: $90 \times 10^6$ cells/mL; high VCD target: $150 \times 10^6$ cells/mL. (A, B) Bleed rates (mL/h) for CLA and CLB perfusion bioreactors, respectively. The bleed allows VCD control in the bioreactors. (C) Cell line B viable cell density (VCD, $\times 10^6$ cells/mL, left y-axis) and viability (%, right y-axis) over the culture period. (D) CLB viability (%) on shortened y-axis scale to highlight the decline in viability at end of culture in high VCD bioreactors. (E, F) Culture glucose concentration (g/L) for CLA and CLB, respectively. (G, H) Culture lactate concentration (g/L) for CLA and CLB, respectively. Two biological replicates were performed per condition and one biological replicate for CLB high VCD.

### Supplementary Figure S5

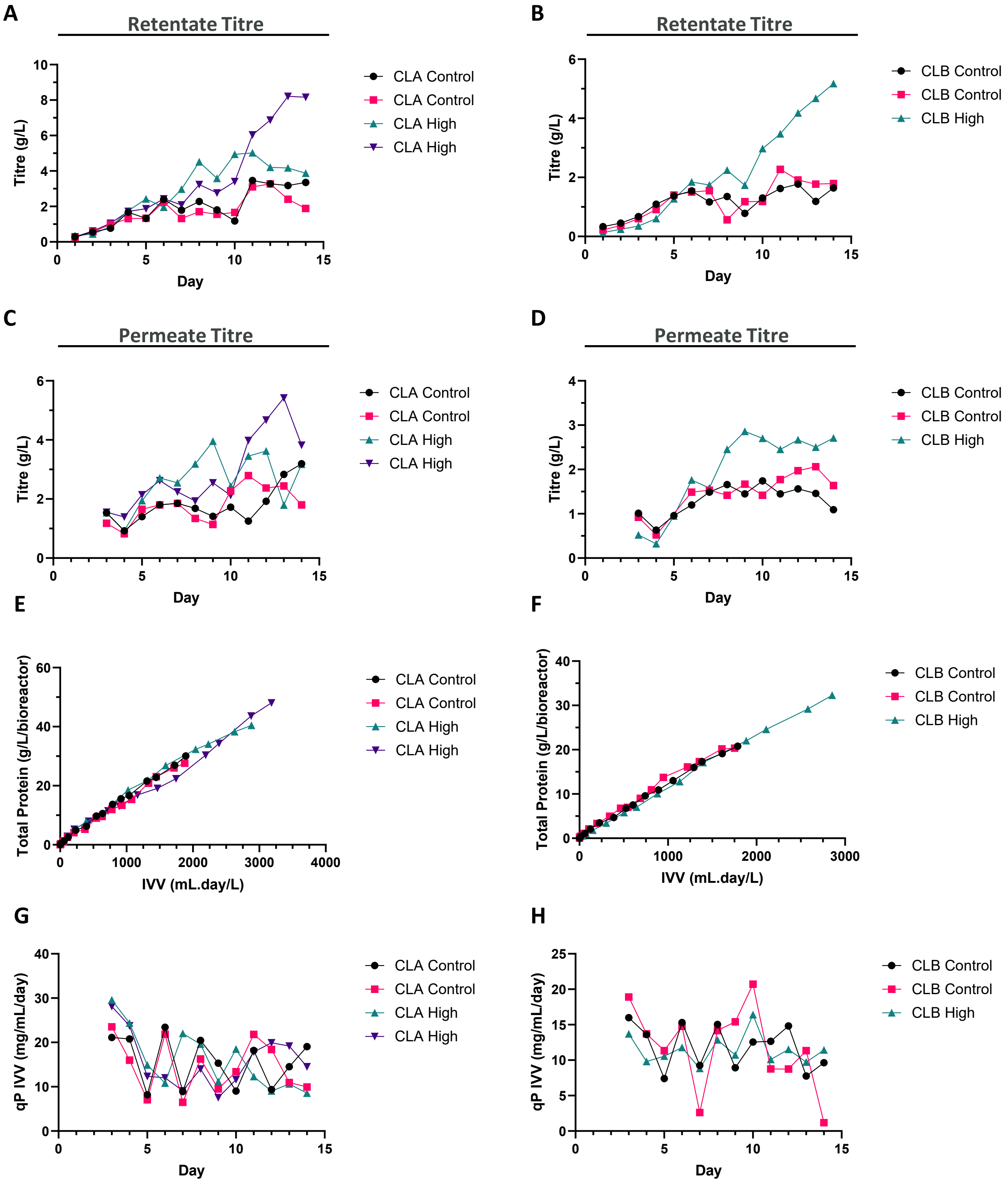

**Supplementary Figure S5. Productivity metrics for Cell line A (CLA) and Cell line B (CLB) perfusion bioreactors.** Bioreactors were operated at control viable cell density (VCD, $90\times10^6$ cells/mL) or high VCD ($150\times10^6$ cells/mL) for 14 days. (A, B) Retentate titre (g/L), antibody concentration retained within the bioreactor, for CLA and CLB, respectively. (C, D) Permeate titre (g/L), antibody concentration in the filtered harvest stream, for CLA and CLB, respectively. (E, F) Cumulative antibody produced versus integral viable volume (IVV, mL·day/L) for CLA and CLB, used to calculate specific cell productivity ($qP_{IVV}$). (G, H) Specific cell productivity ($qP_{IVV}$, mg/mL/day) calculated from the slope of cumulative production versus IVV for CLA and CLB, respectively. Two biological replicates were performed per condition and one biological replicate for CLB high VCD.

#### Supplementary Figure S6

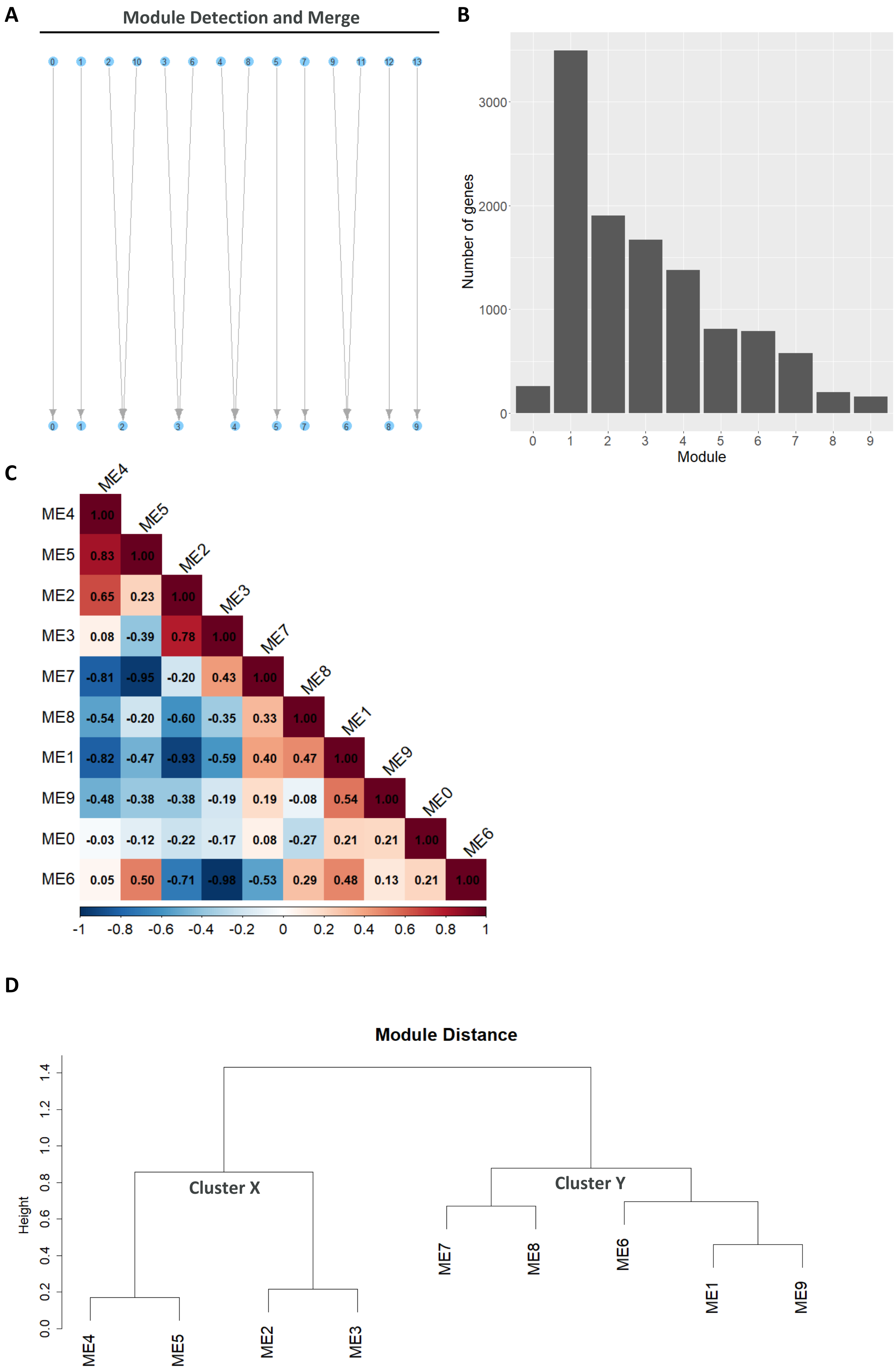

#### Supplementary Figure S6

E

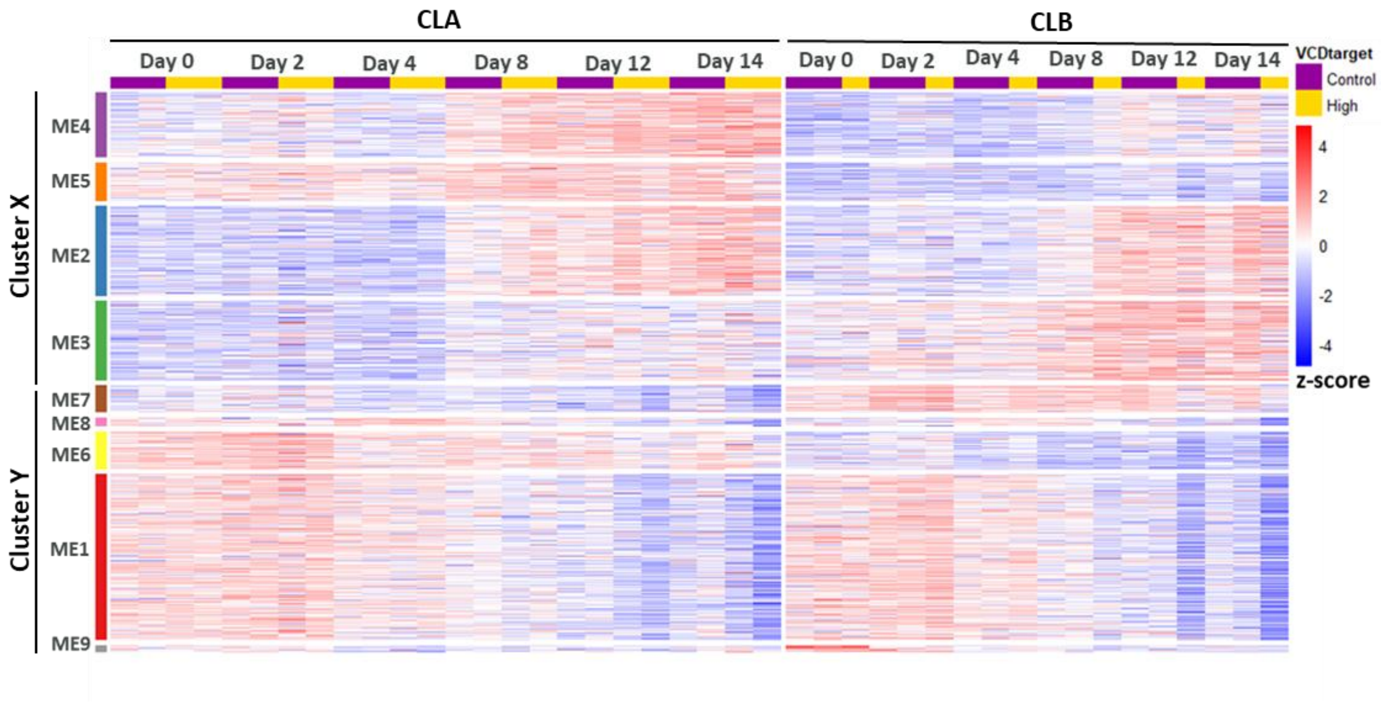


F

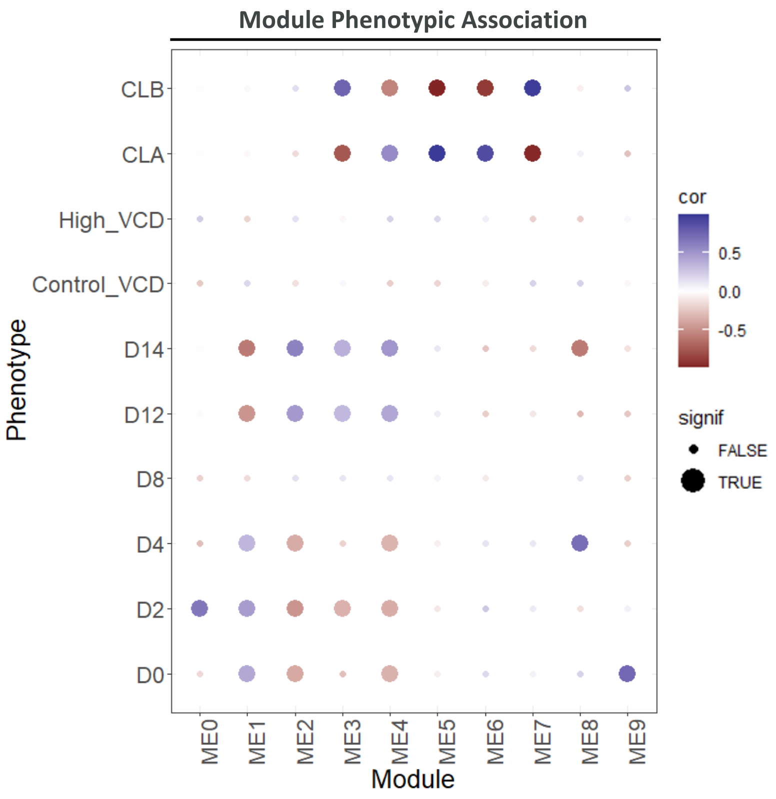

**Supplementary Figure S6. Weighted gene co-expression network analysis (WGCNA) module detection and phenotypic association**. Analysis performed on RNA-seq data normalised counts from Cell line A (CLA) and Cell line B (CLB) perfusion bioreactors sampled on days (D) 0, 2, 4, 8, 12, and 14. (A) Bipartite graph showing initial module detection (14 modules) and subsequent merging of highly similar modules (10 modules), using parameters specified in Materials and Methods. Module 0 was not included as this contained genes which did not cluster into any co-expression module. (B) Number of genes assigned to each of the final modules. (C) Module (ME) eigengene correlation heatmap. Spearman's rank correlation coefficient calculations displaying pairwise correlation between module eigengenes. Colour intensity represents correlation strength, ranging from negative correlations (blue) to positive correlations (red). Values in the heatmap show Spearman's correlation coefficients. (D) ME Spearman distance dendrogram showing hierarchical clustering of the relationships between ME eigengenes based on their distance metrics. Height represents the dissimilarity between ME, with closely related MEs clustering at lower heights, whilst more distinct modules separate at greater heights. Two broad distinct clusters were formed and have been termed cluster X and cluster Y. (E) ME gene expression heatmap over time in the perfusion bioreactor with relevant cluster from ME Spearman's distance labelled. Z-score gene expression is shown. (F) ME-trait correlation dot plot showing phenotypic association of each ME eigengene with culture day, VCD target (90 or $150 \times 10^6$ cells/mL), and cell line (CLA or CLB). Correlation calculated using Pearson's correlation coefficient; blue indicates positive correlation; red indicates negative correlation. Large dots represent a p-value $< 0.05$ indicating statistical significance (signif). Small dots represent a p-value $> 0.05$.

**Supplementary Figure S7**

**A**

| | Likelihood ratio test (LRT) | VCDtarget |
|---|---|---|
| **Full model** | ~Cell line + VCDtarget | ~VCDtarget |
| **Reduced model** | ~Cell line | n/a |

**B**

| | Day 0 | | Day 2 | | Day 4 | | Day 8 | | Day 12 | | Day 14 | |
|---|---|---|---|---|---|---|---|---|---|---|---|---|
| | **LRT** | **VCDtarget** | **LRT** | **VCDtarget** | **LRT** | **VCDtarget** | **LRT** | **VCDtarget** | **LRT** | **VCDtarget** | **LRT** | **VCDtarget** |
| **Upregulated** | 1 | 0 | 0 | 0 | 0 | 0 | 26 | 8 | 63 | 17 | 100 | 30 |
| **Downregulated** | 0 | 0 | 0 | 0 | 0 | 0 | 193 | 79 | 256 | 99 | 246 | 108 |

**Supplementary Figure S7. Differential gene expression analysis comparing high (150x$10^6$ cells/mL) vs control (90x$10^6$ cells/mL) viable cell density (VCD) bioreactors on each day independently using two design models**. (A) Summary of the two design models: the likelihood ratio test (LRT), which tests whether adding VCD target significantly improves model fit for each gene, and the VCD target model, which excludes cell line as a confounding variable. (B) Number of differentially expressed genes (DEGs) from both models. Overall, the LRT model detects more DEGs than the VCD target model.

#### Supplementary Figure S8

**A**

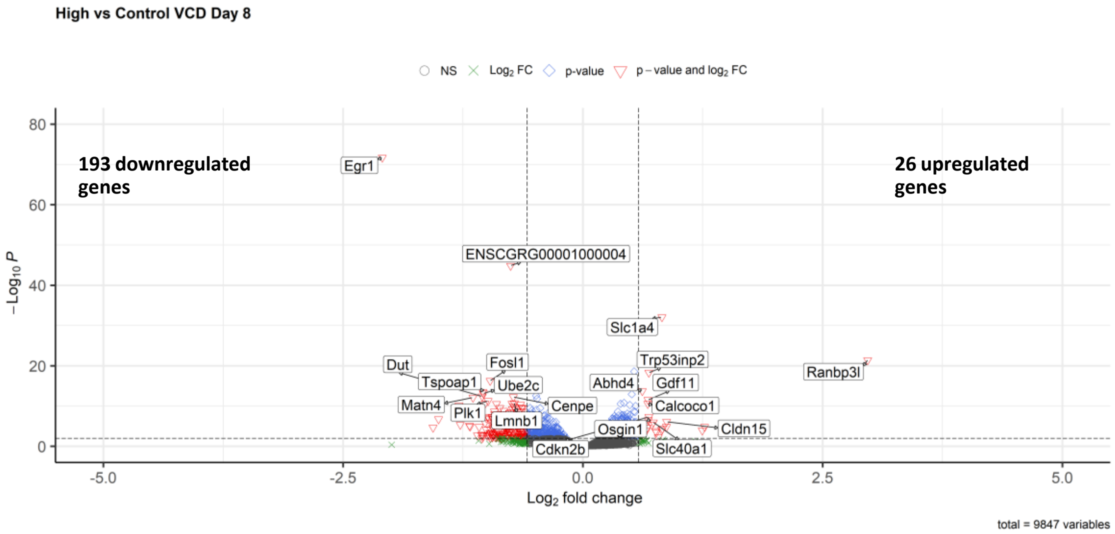


**B**

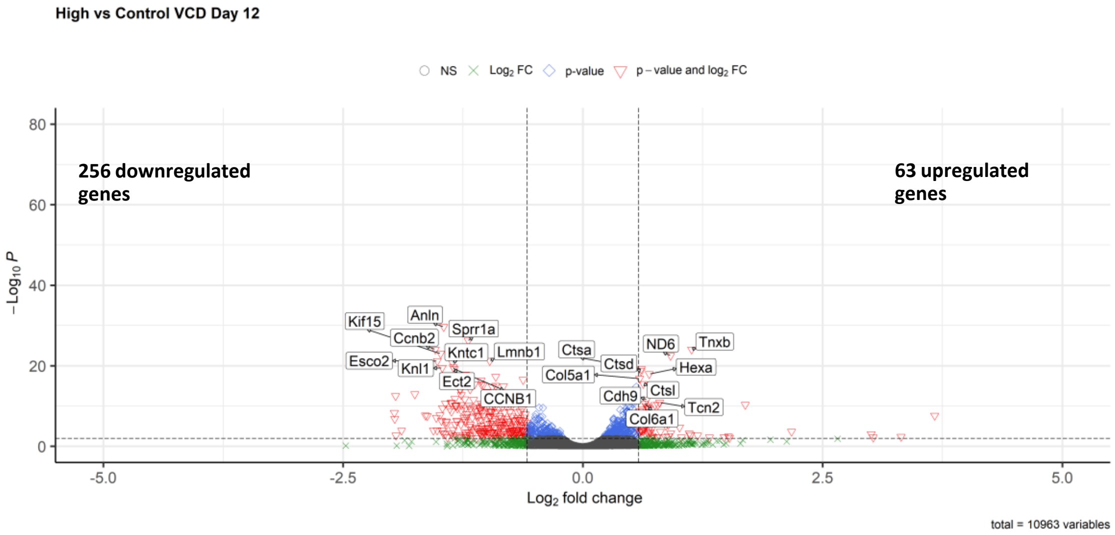


**C**

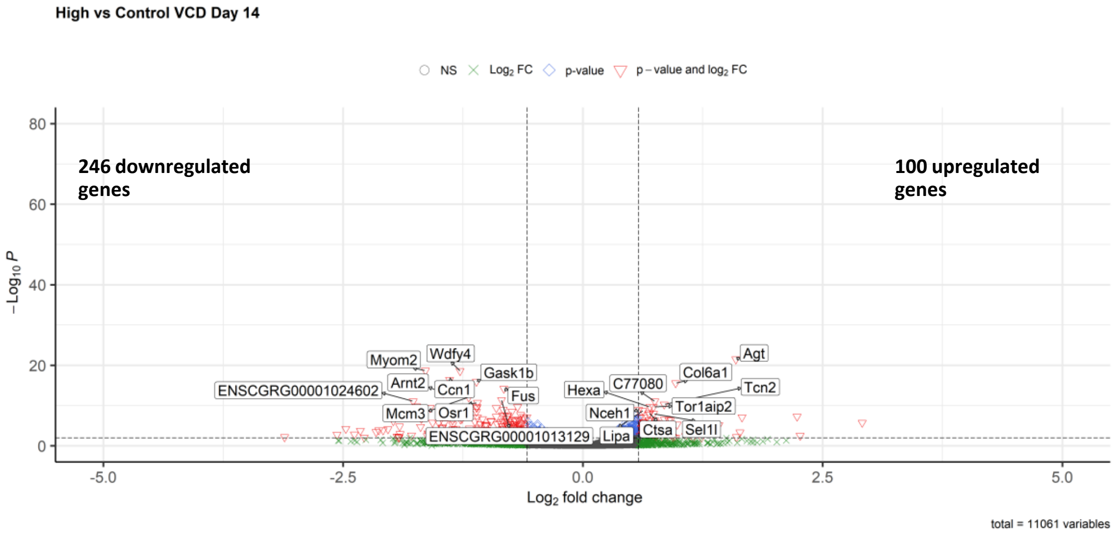

**Supplementary Figure S8. Differential gene expression analysis comparing high viable cell density (VCD) to control VCD bioreactors each day independently controlling for cell line variation (Likelihood ratio test model, LRT).** Volcano plots showing $\log_2$ fold change against $-\log_{10}$(adjusted p-value) for (A) day 8, (B) day 12, and (C) day 14. High VCD: 150x$10^6$ cells/mL; control VCD: 90x$10^6$ cells/mL. Differential gene expression analysis performed using DESeq2 with clone included as a covariate to control for cell line differences. Significantly differentially expressed genes (DEGs) defined as adjusted p-value <0.01 and $\log_2$ fold change >0.58 (equivalent to >1.5-fold change). The top 10 up and downregulated differentially expressed genes are labelled. The total number of up and downregulated genes meeting the DEG criteria are situated in top right and top left, respectively. The number of genes tested is situated on the bottom right of each volcano plot. Adjusted p-values calculated using Benjamini-Hochberg false discovery rate (FDR) correction.

**Supplementary Figure S9**

A A

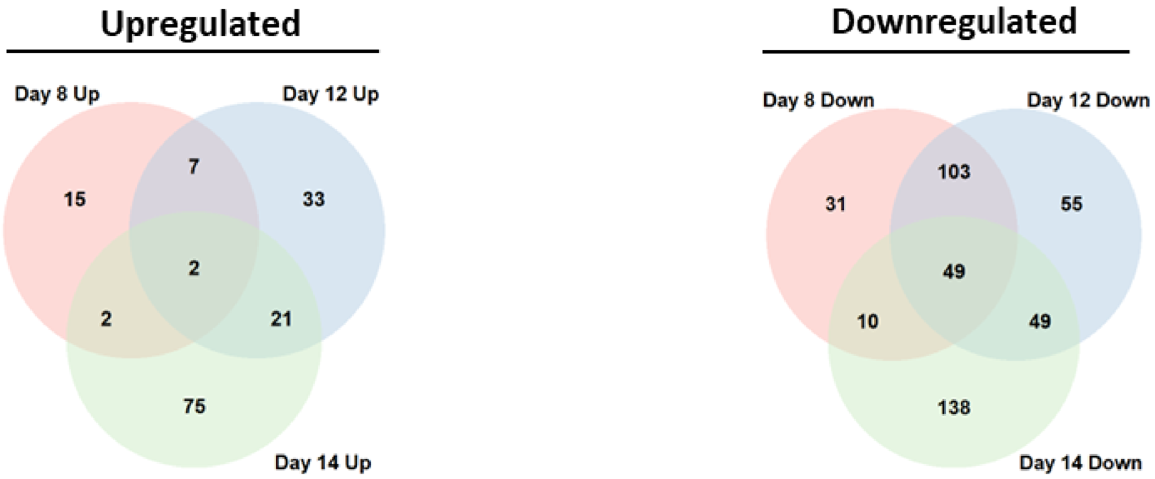


**Supplementary Figure S9. Overlap of differentially expressed genes (DEGs) on days 8, 12 and 14 from comparisons of high viable cell density (VCD) to control VCD bioreactors each day independently controlling for cell line variation (Likelihood ratio test model, LRT).** Significant DEGs defined as having an adjusted p-value <0.01 and $log_2$ fold change >0.58 (equivalent to >1.5-fold change). (A) Upregulated gene overlap. (B) Downregulated gene overlap.

### Supplementary Figure S10

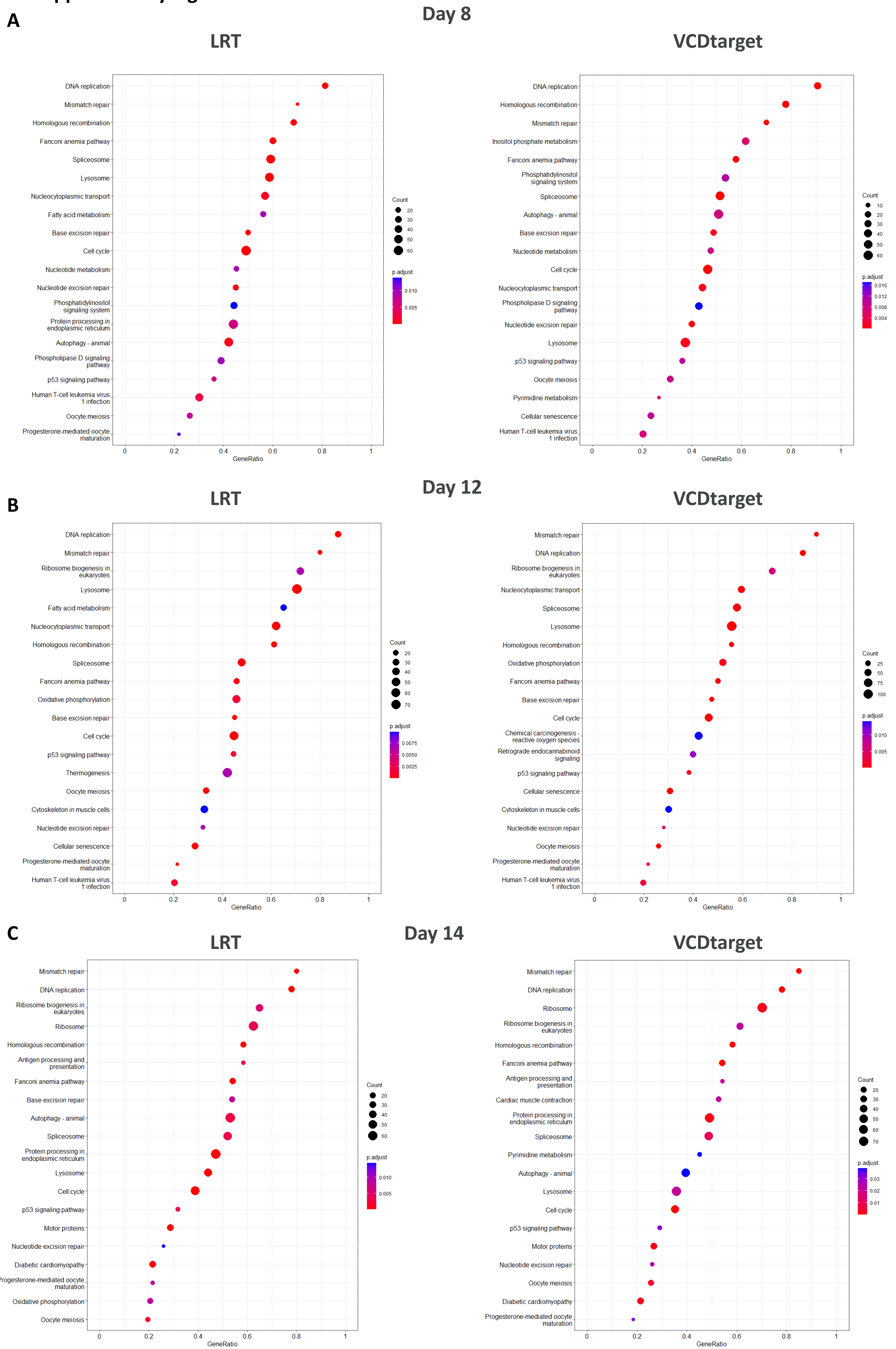

**Supplementary Figure S10. Differential gene expression analysis comparing high ($150x10^6$ cells/mL) vs control ($90x10^6$ cells/mL) viable cell density (VCD) bioreactors on each day independently using two design models genes gene set enrichment analysis (GSEA).** (A), (B), (C), KEGG pathway gene set enrichment analysis (GSEA) comparing high to control VCD bioreactors on (A) Day 8, (B), Day 12 and (C) Day 14 contrasting results from the likelihood ratio test (LRT) model and the model omitting cell line variation from the design (VCDtarget). Dot plots showing enriched KEGG pathways performed using clusterProfiler with genes ranked by $log_2$ fold change. Dot size represents gene ratio (proportion of genes in pathway that are differentially expressed); colour indicates adjusted p-value (Benjamini-Hochberg FDR correction). Enrichment analysis between both models were highly similar.

**Supplementary Figure S11**

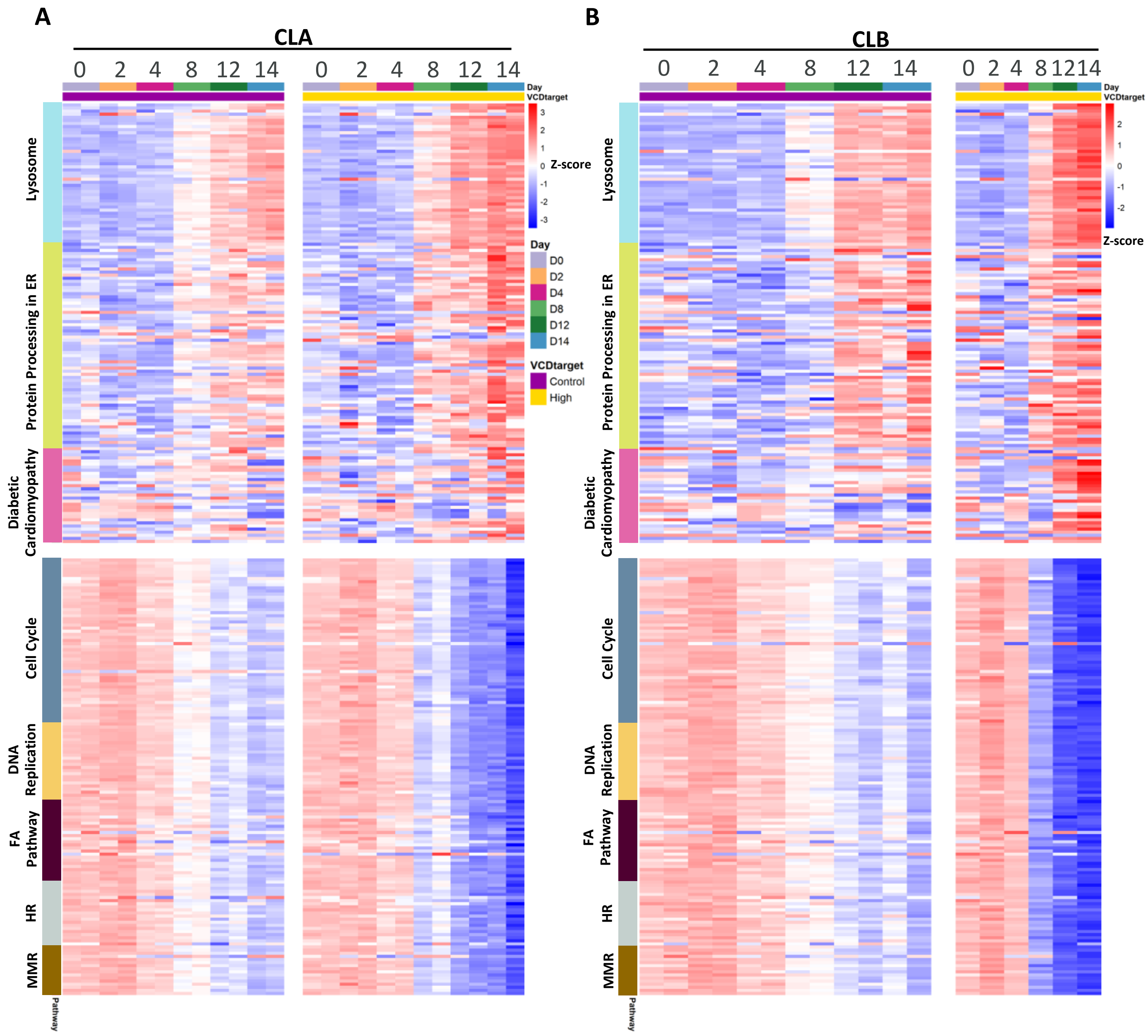


**Supplementary Figure S11. KEGG gene set enrichment analysis (GSEA) pathway gene expression.** (A, B) Heatmaps showing z-score normalised expression of genes within significantly enriched KEGG pathways for (A) Cell line A (CLA) and (B) Cell line B (CLB). Samples are grouped by VCD condition (high: 150x10$^6$ cells/mL; control: 90x10$^6$ cells/mL), and pathways are separated into upregulated and downregulated categories.

**Supplementary Figure S12**

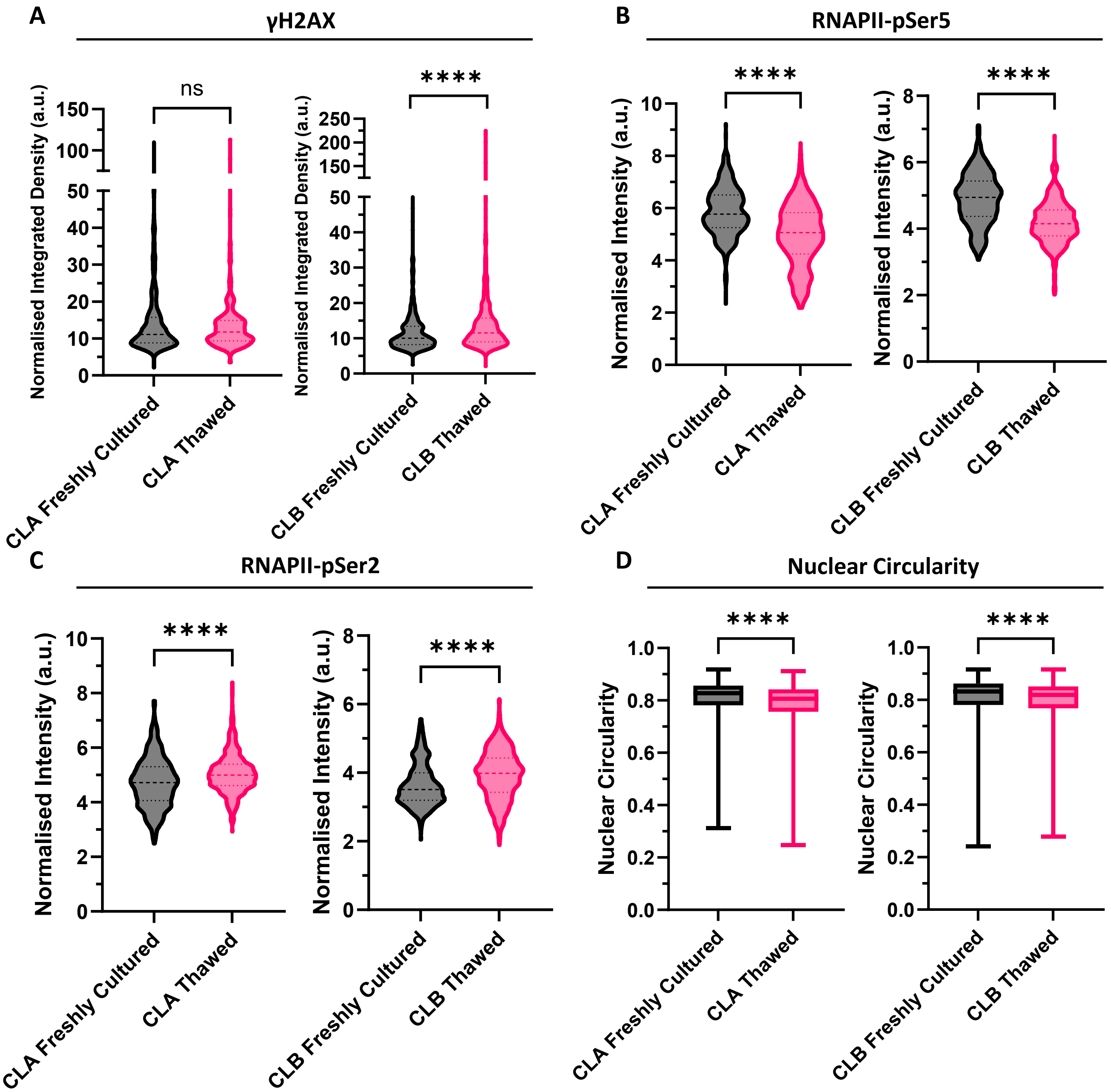


**Supplementary Figure S12. Investigating the impact of thaw-induced effects in CHO cells.** (A) Quantification of γH2AX integrated density normalised to background fluorescence. Cell line A (CLA) freshly cultured: n = 504, CLA thawed: n = 530, Cell line B (CLB) freshly cultured: 550, CLB thawed: n = 505. (B) Quantification of RNAPII-pSer5 fluorescence intensity normalised to background fluorescence. CLA freshly cultured: n = 519, CLA thawed: n = 541, CLB freshly cultured: 533, CLB thawed: n = 535. (C) Quantification of RNAPII-pSer2 fluorescence intensity normalised to background fluorescence. CLA freshly cultured: n = 512, CLA thawed: n = 527, CLB freshly cultured: 557, CLB thawed: n = 543. (D) Nuclear circularity measurements. CLA freshly cultured: n = 1535, CLA thawed: n = 1598, CLB freshly cultured: 1640, CLB thawed: n = 1583. All statistics determined by Mann-Whitney U test. Three biological replicates were performed per condition. Significance definitions, ns $p > 0.05$, * $p < 0.05$, **** $p < 0.0001$.

#### Supplementary Figure S13

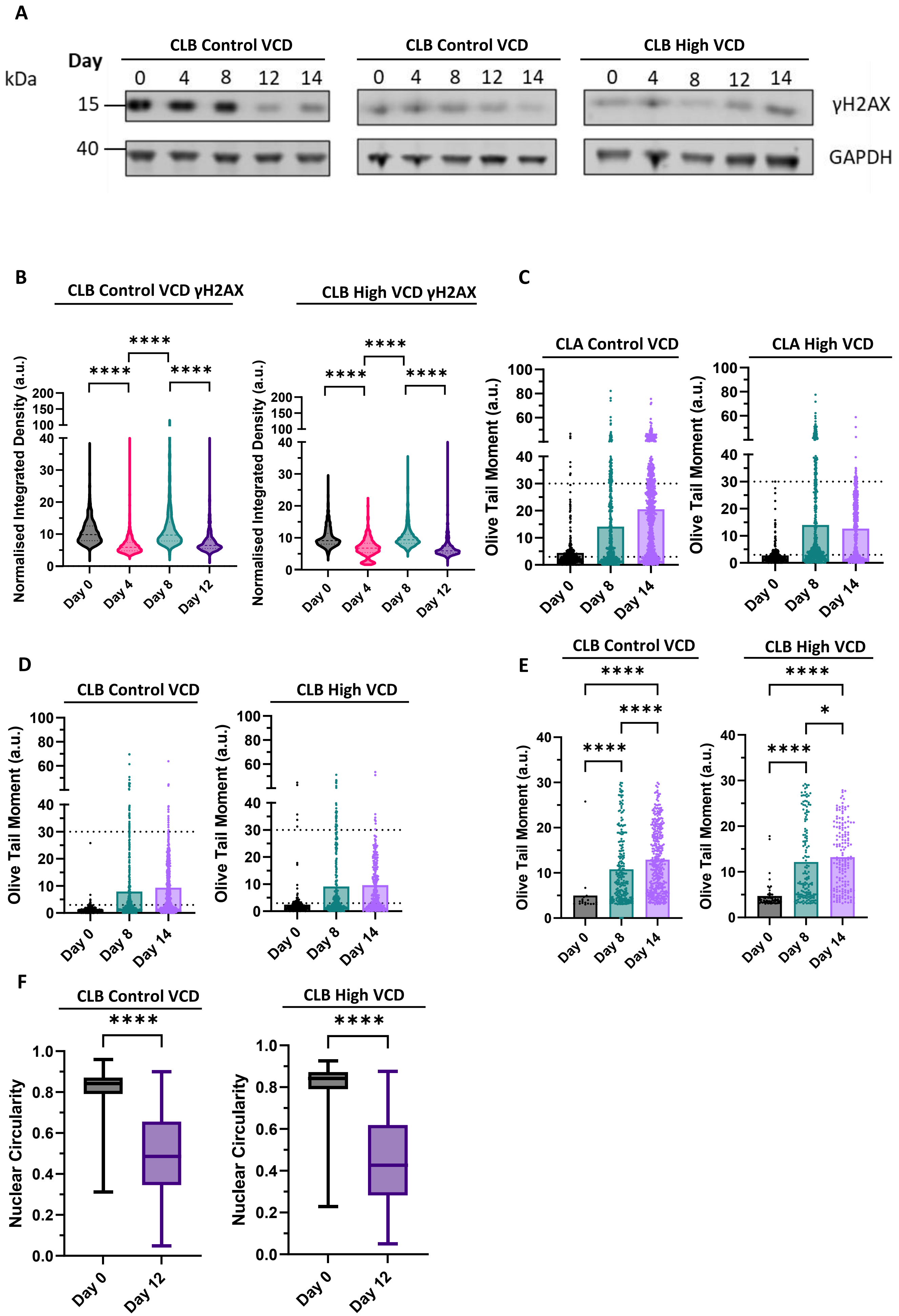

**Supplementary Figure S13. Assessment of DNA damage and nuclear architecture.** (A) Representative γH2AX western blots for Cell line B (CLB) control and high viable cell density (VCD) bioreactors. (B) Quantification of γH2AX integrated density normalised to background fluorescence. CLB control VCD: n = 1024/1046/1034/1118, CLB high VCD: n = 533/554/508/553, statistics determined by Kruskal-Wallis test with Dunn's multiple comparisons test. (C) Cell line A (CLA) and (D) CLB full alkaline comet assay quantification analysis. CLA control VCD: n = 439/659/817, CLA high VCD: n = 405/551/854, CLB control VCD: n = 424/662/801, CLB high VCD: n = 332/362/275. (E) CLB control and high VCD DNA damaged subpopulation alkaline comet assay. CLB control VCD: n = 18/236/380, CLB high VCD: n = 53/147/164, statistics determined by Kruskal-Wallis test with BKY multiple comparisons test. (F) Quantification of nuclear circularity, CLB control VCD: n = 3110/3240, CLB high VCD: n = 1591/1647, statistics determined by Mann-Whitney U test. Two biological replicates were performed per condition and one biological replicate for CLB high VCD. Significance definitions, ns $p > 0.05$, *$p < 0.05$, ****$p < 0.0001$.

#### Supplementary Figure S14

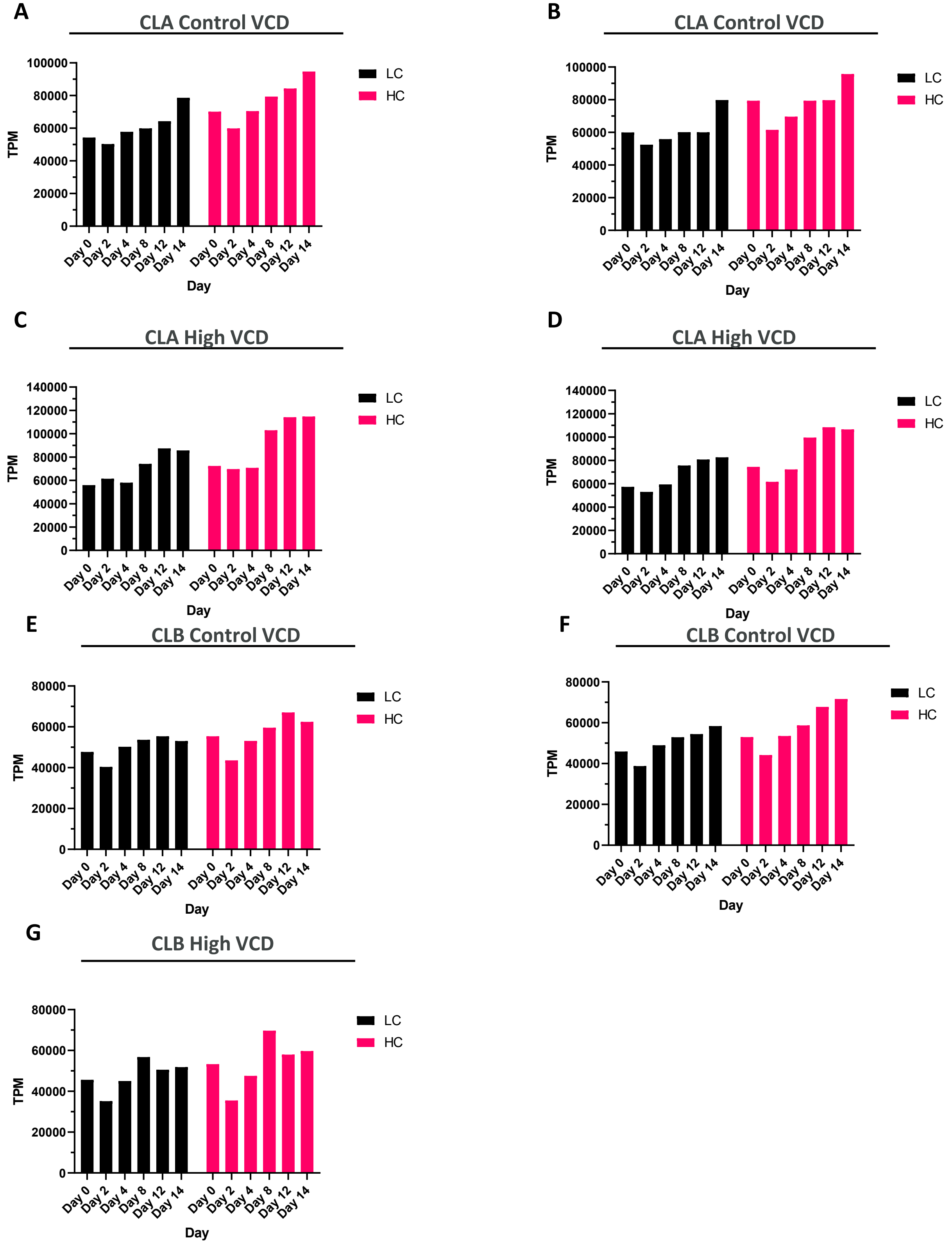


**Supplementary Figure S14. Heavy (HC) and light chain (LC) transcript levels during perfusion bioreactor culture.** Transcript abundance measured as transcripts per million (TPM) from RNA-seq data over time (days 0, 2, 4, 8, 12, 14). (A) Cell line A (CLA) control viable cell density (VCD), (B) CLA control VCD, (C) CLA high VCD, (D) CLA high VCD, (E) Cell line B (CLB) control VCD, (F) CLB control VCD, (G) CLB high VCD.

**Supplementary Figure S15**

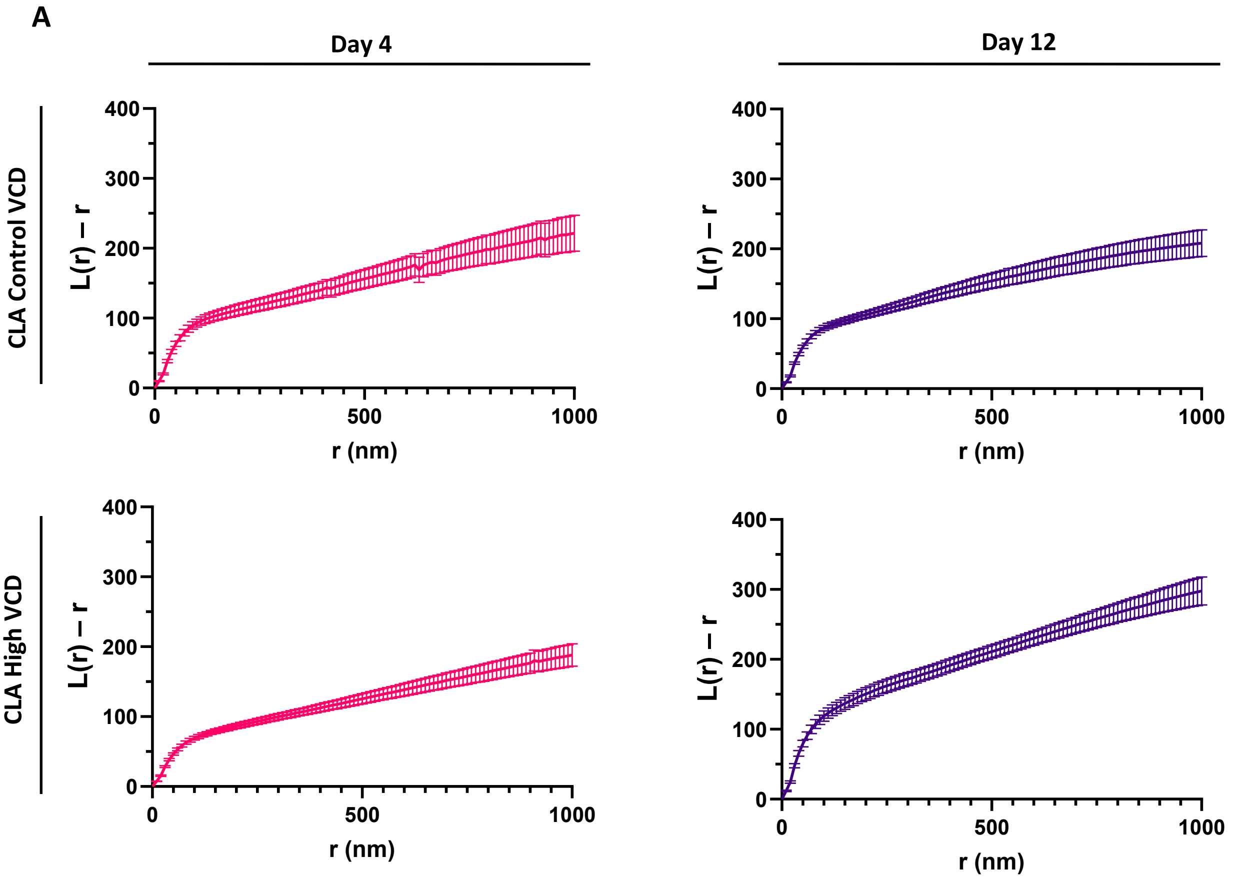


**Supplementary Figure S15. Spatial clustering analysis of RNA polymerase II (RNAPII)-pSer5 using linearised Ripley's K function.** (A) Cell line A (CLA) control viable cell density (VCD, $90x10^6$ cells/mL) and high VCD ($150x10^6$ cells/mL) at days 4 and 12. Linearised Ripley's K function, L(r) - r, calculated from STORM super-resolution microscopy images, where r is the spatial radius (nm). L(r) - r = 0 indicates random spatial distribution; positive values indicate molecular clustering. Data represent mean values ± SD. CLA control VCD: n = 31/29, CLA high VCD: n = 36/31. One biological replicate performed per condition.
