## Supplementary tables for "DNA Damage Driven Viability Loss and Transcriptional Reprogramming in Chinese Hamster Ovary Cell Perfusion Culture"

### Supplementary Table 1

| Target | Supplier | Catalogue Number | Assay |
|---|---|---|---|
| Phospho-Histone H2A.X (Ser139) (20E3) | Cell Signaling Technology | 9718 | Western blot (1:1000)/Immunofluorescence (1:200) |
| GAPDH (14CLA0) | Cell Signaling Technology | 2118 | Western blot (1:1000) |
| GAPDH (D4C6R) | Cell Signaling Technology | 97166 | Western blot (1:1000) |
| Anti-RNA polymerase II CTD repeat YSPTSPS (phospho S5) antibody [4H8] | Abcam | Ab5408 | Western blot (1:1000) |
| Anti-RNA polymerase II CTD repeat YSPTSPS (phospho S2) | Abcam | ab5095 | Western blot (1:1000) |
| Rpb1 CTD (4H8) | Cell Signaling Technology | 2629 | Western blot (1:1000) |
| Alexa Fluor® 488 Anti-RNA polymerase II CTD repeat YSPTSPS (phospho S5) | Abcam | ab196152 | Immunofluorescence (1:500)/STORM (1:500) |
| Alexa Fluor® 488 Anti-RNA polymerase II CTD repeat YSPTSPS (phospho S2) | Abcam | ab237279 | Immunofluorescence (1:200) |
| Donkey F(ab')2 Anti-Rabbit IgG H&L (Alexa Fluor® 488) preadsorbed | Abcam | ab181346 | Immunofluorescence (1:500) |
| IRDye 800 CW Goat anti-Rabbit IgG Antibody | LI-COR bio | 926-32211 | Western blot (1:10,000) |
| IRDye 680RD Goat anti-Mouse IgG Antibody | LI-COR bio | 926-68070 | Western blot (1:10,000) |

**Supplementary Table 1. Antibodies used for western blot, immunofluorescence, and STORM super-resolution microscopy.** Table lists primary and secondary antibodies including target protein, host species, clonality, supplier, catalogue number, and dilutions used for each application.

### Supplementary Table 2

| Post-Translational Modification | CLA Control Day 5 rep 1 (%) | CLA Control Day 14 rep 1 (%) | CLA Control Day 5 rep 2 (%) | CLA Control Day 14 rep 2 (%) | CLA High Day 5 rep 1 (%) | CLA High Day 14 rep 1 (%) | CLA High Day 5 rep 2 (%) | CLA High Day 14 rep 2(%) | CLB Control Day 5 rep 1(%) | CLB Control Day 14 rep 1 (%) | CLB Control Day 5 rep 2 (%) | CLB Control Day 14 rep 2 (%) | CLB High Day 5 rep 1 (%) | CLB High Day 14 rep 1 (%) |
|---|---|---|---|---|---|---|---|---|---|---|---|---|---|---|
| **HC M256 oxidation** | 2.5 | 2.4 | 2.6 | 1.9 | 2.6 | 1.8 | 2.5 | 1.7 | 1.8 | 1.8 | 2.2 | 1.8 | 2.3 | 2 |
| **HC N329 succinimide** | 2 | 5.6 | 2.2 | 5.7 | 2.9 | 6.1 | 3.2 | 6.7 | 3.7 | 7 | 4 | 7.3 | 4.9 | 7.3 |
| **HC N388/N393/N394 succinimide** | 1.9 | 2 | 2 | 2.1 | 2 | 2 | 2 | 2 | 2.1 | 2 | 2 | 2.1 | 2.1 | 2 |
| **HC W421/M432 oxidation** | 1.3 | 1.4 | 1.3 | 1.1 | 1.4 | 1 | 1.3 | 1 | 1 | 1.1 | 1.2 | 1.1 | 1.3 | 1.1 |
| **LC Q6 deamidation** | 1.1 | 1.5 | 1 | 1 | 1.2 | <LOQ | 1.1 | <LOQ | <LOQ | <LOQ | <LOQ | 1 | 1.3 | 1 |
| **LC VHS leader seq** | 2.6 | 3.3 | 2.5 | 3.3 | 2.7 | 3.5 | 2.6 | 3.1 | 2.8 | 2.4 | 3.2 | 2.3 | 3.2 | 2.4 |

**Supplementary Table 2. Post-translational modifications (PTMs) in mAb product assessed by tryptic peptide mapping LC-MS/MS.** PTM analysis of purified antibody from Cell line A (CLA) and Cell line B (CLB) perfusion bioreactors at days 5 and 14, comparing control viable cell density (VCD, $90x10^6$ cells/mL) and high VCD ($150x10^6$ cells/mL) conditions. Table shows relative abundance (%) of major PTMs. Sample preparation and analysis performed as described in materials and methods. Two biological replicates were performed per condition and one biological replicate for CLB high VCD.

### Supplementary Table 3

| Sequence Variants | Ab Chain | CLA Control Day 5 rep 1 (%) | CLA Control Day 14 rep 1 (%) | CLA Control Day 5 rep 2 (%) | CLA Control Day 14 rep 2 (%) | CLA High Day 5 rep 1 (%) | CLA High Day 14 rep 1 (%) | CLA High Day 5 rep 2 (%) | CLA High Day 14 rep 2(%) | CLB Control Day 5 rep 1(%) | CLB Control Day 14 rep 1 (%) | CLB Control Day 5 rep 2 (%) | CLB Control Day 14 rep 2 (%) | CLB High Day 5 rep 1 (%) | CLB High Day 14 rep 1 (%) |
|---|---|---|---|---|---|---|---|---|---|---|---|---|---|---|---|
| **A122S** | HC | 0.3 | 0.4 | 0.3 | 0.3 | 0.3 | 0.4 | 0.3 | 0.7 | 0.1 | 0.6 | 0.2 | 0.8 | 0.1 | 0.3 |
| **I48R** | LC | 0.5 | 0.5 | 0.4 | 0.6 | 0.5 | 0.5 | 0.5 | 0.5 | 0.5 | 0.5 | 0.5 | 0.6 | 0.6 | 0.6 |
| **L116R** | HC | 0.3 | 0.4 | 0.3 | 0.4 | 0.3 | 0.3 | 0.3 | 0.3 | 0.3 | 0.3 | 0.3 | 0.3 | 0.3 | 0.3 |
| **L47R** | LC | 0.5 | 0.5 | 0.4 | 0.6 | 0.5 | 0.5 | 0.5 | 0.5 | 0.5 | 0.5 | 0.5 | 0.6 | 0.6 | 0.6 |
| **P221L** | HC | 41.4 | 37.6 | 43 | 37.4 | 41 | 37 | 40.7 | 38.1 | 40.5 | 39.2 | 43.8 | 38.9 | 38.6 | 37.9 |

**Supplementary Table 3. Sequence variants in mAb product assessed by tryptic peptide mapping LC-MS/MS at days 5 and 14.** Analysis of amino acid sequence variants in purified antibody from Cell line A (CLA) and Cell line B (CLB) perfusion bioreactors, comparing control viable cell density (VCD, $90x10^6$ cells/mL) and high VCD ($150x10^6$ cells/mL) conditions. Table shows relative abundance (%) of detected sequence variants. Note: P221L variant abundance is overestimated due to the short tryptic peptide containing this position. Sample preparation and analysis performed as described in materials and methods. Two biological replicates were performed per condition and one biological replicate for CLB high VCD.

**Supplementary Table 4**

| Sample | %LMWS |
|---|---|
| CLA Control Day 5 rep 1 | 2.4 |
| CLA Control Day 14 rep 1 | 3.7 |
| CLA Control Day 5 rep 2 | 2.8 |
| CLA Control Day 14 rep 2 | 3.6 |
| CLA High Day 5 rep 1 | 3.6 |
| CLA High Day 14 rep 1 | 3.5 |
| CLA High Day 5 rep 2 | 3.2 |
| CLA High Day 14 rep 2 | 3.5 |
| CLB Control Day 5 rep 1 | 3.3 |
| CLB Control Day 14 rep 1 | 3.5 |
| CLB Control Day 5 rep 2 | 2.9 |
| CLB Control Day 14 rep 2 | 3.4 |
| CLB High Day 5 rep 1 | 3.1 |
| CLB High Day 14 rep 1 | 3.3 |

**Supplementary Table 4: Low molecular weight species (LMWS) analysis at days 5 and 14 by microchip capillary electrophoresis (MCE).** LMW species quantified using the LabChip GXII Touch instrument under non-reducing conditions. Purified mAb samples from Cell line A (CLA) and Cell line B (CLB) perfusion bioreactors at control viable cell density (VCD, 90x$10^6$ cells/mL) and high VCD (150x$10^6$ cells/mL) were analysed to determine the percentage of LMW species (fragments) relative to intact antibody. Sample preparation and analysis performed as described in materials and methods. Two biological replicates were performed per condition and one biological replicate for CLB high VCD.
